# Cryo-EM structure of the mitochondrial protein-import channel TOM complex from *Saccharomyces cerevisiae*

**DOI:** 10.1101/798371

**Authors:** Kyle Tucker, Eunyong Park

## Abstract

Nearly all mitochondrial proteins are encoded by the nuclear genome and imported into mitochondria following synthesis on cytosolic ribosomes. These precursor proteins are translocated into mitochondria by the TOM complex, a protein-conducting channel in the mitochondrial outer membrane. Using cryo-EM, we have obtained high-resolution structures of both apo and presequence-bound core TOM complexes from *Saccharomyces cerevisiae* in dimeric and tetrameric forms. Dimeric TOM consists of two copies each of five proteins arranged in two-fold symmetry—Tom40, a pore-forming β-barrel with an overall negatively-charged inner surface, and four auxiliary α-helical transmembrane proteins. The structure suggests that presequences for mitochondrial targeting insert into the Tom40 channel mainly by electrostatic and polar interactions. The tetrameric complex is essentially a dimer of dimeric TOM, which may be capable of forming higher-order oligomers. Our study reveals the molecular organization of the TOM complex and provides new insights about the mechanism of protein translocation into mitochondria.

## Introduction

Mitochondria are double-membrane-bound organelles that perform oxidative phosphorylation and other essential cellular functions in eukaryotic cells. There are ∼1,000–1,500 mitochondrial proteins and the vast majority (∼99%) are synthesized by cytosolic ribosomes, initially as precursor proteins that are then imported into mitochondria^1-3^. Multiple protein complexes within the organelle mediate membrane translocation and sorting of these precursor polypeptides into four distinct compartments—the outer membrane, the inner membrane, the intermembrane space (IMS), and the matrix. The general import pore in the outer membrane is formed by the TOM complex (*T*ranslocase of the *O*uter *M*embrane), which is responsible for initial translocation of over 90% of mitochondrial precursor proteins from the cytosol to the IMS.

Studies of the TOM complex of fungal cells have established that it consists of seven transmembrane proteins: Tom40, Tom22, Tom5, Tom6, and Tom7, as well as Tom70 and Tom20 (ref. 4,5). The first five proteins form a stable complex, referred to as the core TOM complex, whereas the latter two readily dissociate from the core complex upon isolation in detergent^6,7^. Various analyses have indicated that the detergent-solubilized TOM complex has an apparent molecular mass of ∼400–600 kDa and contains multiple copies of each Tom subunit^6-10^. The translocation pore through which precursor polypeptides must pass is formed by Tom40 (ref. 5,11-13), a β-barrel protein structurally related to the voltage-dependent anion-selective channel VDAC, a major mitochondrial outer membrane porin^14,15^. The other Tom proteins are associated with Tom40 by their single α-helical transmembrane segments (TMs). Although functions of the α-helical Tom subunits are relatively poorly defined, they have been suggested to act either as receptors for precursor proteins^16-20^ or as binding sites for other factors^20,21^, and/or as escorts that promote assembly and stability of the TOM complex^6,10,22,23^.

Current evidence indicates that translocation is a sequential process in which a precursor protein is first recruited by the cytosolic receptor domains of Tom70, Tom20, and Tom22, then threaded into the pore of Tom40, and finally handed over to the translocase of the inner membrane (TIM) complex or IMS-resident chaperones (for review, see ref. 2). However, the underlying mechanism by which the TOM complex enables these events has been unclear. In particular, how the Tom40 channel interacts with mitochondrial targeting sequences is poorly understood. Interaction of such presequences (N-terminal cleavable sequences that target ∼60–70% mitochondrial precursor proteins) is a key step in translocation^11,24-26^. A cryo-electron microscopy (cryo-EM) structure of an apo form of the core TOM complex from *Neurospora crassa* was reported^27^, but its relatively low resolution (∼7-Å) offered only limited insight and precluded building of an atomic model. In addition, the oligomeric architecture of the TOM complex remained a puzzle. The *N. crassa* structure represents a dimeric complex in which two identical pores are symmetrically arranged. However, based on previous low-resolution electron microscopy (EM) and crosslinking analyses, it has been generally thought that the TOM complex is rather dynamic and that the mature form is a trimer^5,13,28,29^. The functional states of the different oligomers remain unclear.

Here we describe three cryo-EM structures of the core TOM complex from *Saccharomyces cerevisiae* that we have determined at near-atomic resolution—an apo and presequence-bound dimeric complex and an apo tetrameric complex. These new structures provide for the first time molecular details of the overall architecture of the TOM complex and, most importantly, the structure of the Tom40 pore and how it engages with presequences.

### Purification and Cryo-EM analysis of the yeast dimeric TOM complex

To enable efficient structural analysis, we first developed a new approach to overexpress and purify the *S. cerevisiae* TOM complex. All Tom subunits, except for Tom70, were expressed in yeast cells from an inducible promoter. We omitted Tom70 because it is known to easily dissociate from the rest of the complex even under very mild purification conditions^9,29^. The complex was directly isolated from whole-cell lysate by tandem affinity purification, utilizing His- and Strep-tags C-terminally attached to Tom22 and Tom40, respectively. The complex was initially extracted with lauryl maltose neopentyl glycol (LMNG) detergent but was exchanged into dodecylmaltoside (DDM) during affinity purification. The TOM complex purified by this method eluted in size-exclusion chromatography (SEC) as a sharp monodisperse peak containing Tom40 and other Tom subunits roughly at stoichiometric ratios (Supplementary Fig. 1a). The purified sample did not contain Tom20 although it was included in overexpression, perhaps because of its low-affinity association with the core complex^6,9^.

To determine the structure of the purified TOM complex, we used single-particle cryo-EM analysis (Supplementary Fig. 1). To gain insight into how TOM recognizes precursor proteins, we added to the purified complex a 23-amino acid-long presequence peptide (pALDH) derived from rat aldehyde dehydrogenase^18,25^, which is predicted to form a positively-charged amphipathic helix like other canonical presequences. We collected ∼460,000 particle images and subjected them to reference-free two-dimensional (2D) classification. Resulting class averages showed that the complex is predominantly a dimer (Supplementary Fig. 1d), closely resembling images reported for the *N. crassa* structure^27^. Both 2D and three-dimensional (3D) classifications indicated high structural homogeneity of the sample (Supplementary Fig. 1c, d). After excluding empty detergent micelle and low-quality particles, ∼70% of particle images (160,577 out of 243,227) were used for the final 3D reconstruction of the dimeric TOM complex at 3.1-Å resolution with C2 symmetry imposed (Fig. 1a, b, Supplementary Figs. 1, 2). Without imposing symmetry, the map was refined to slightly lower resolution (3.2 Å) and manifested no noticeable differences from the symmetrically refined reconstruction (cross-correlation=0.99; data not shown), indicating that the dimer is highly symmetric. To compare structures with or without a bound presequence peptide, we also collected a smaller dataset for the apo TOM complex without the presequence peptide, which produced a 3.5-Å-resolution map (Supplementary Fig. 3). The apo TOM and TOM-pALDH structures are essentially identical (RMSD=∼0.3 Å), except that the latter contains an additional feature in the Tom40 pores corresponding to the presequence peptide (Supplementary Fig 3g; see below).

**Figure 1.**
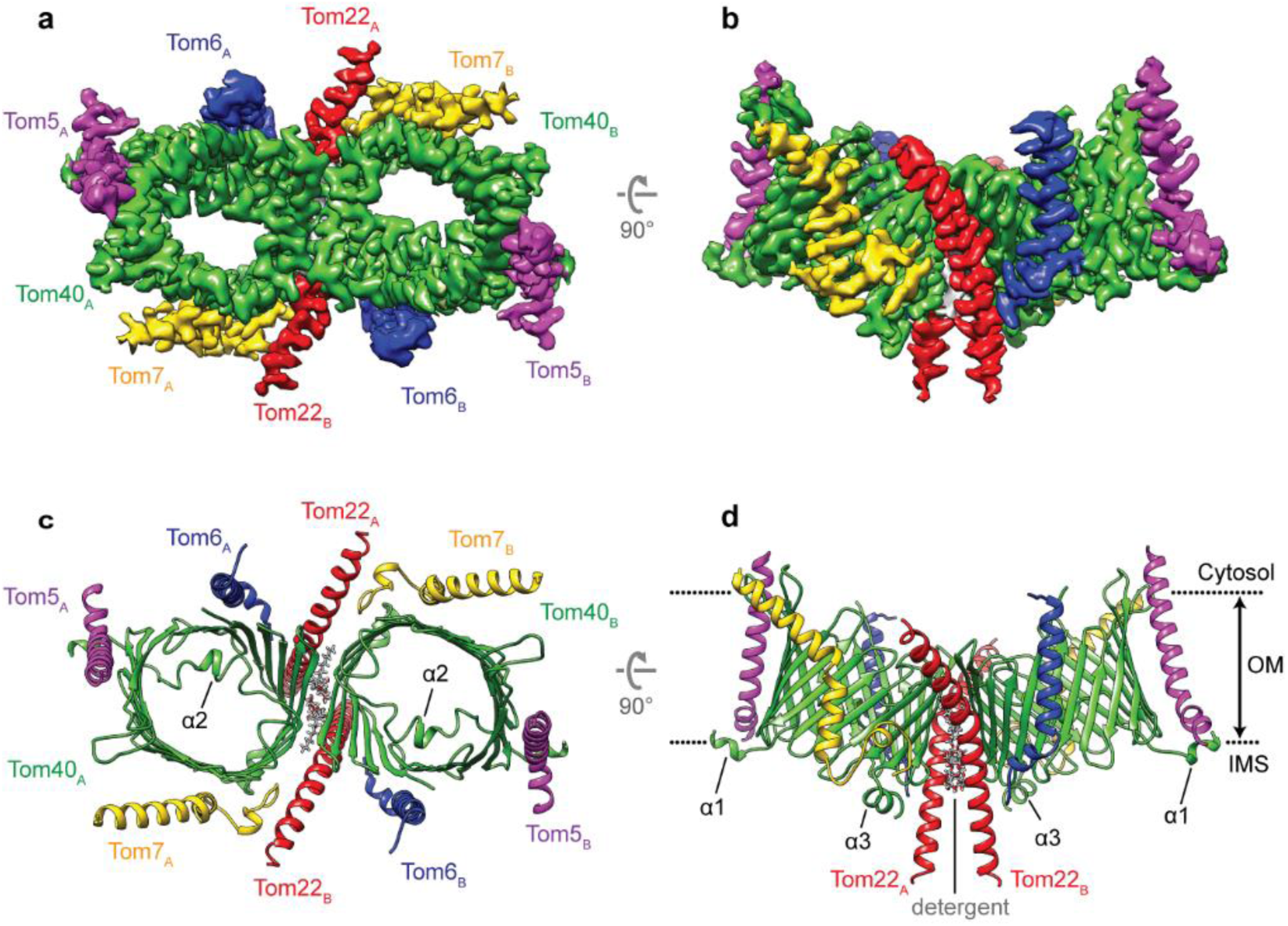
Structure of the dimeric core TOM complex from *S. cerevisiae*. **a, b**, 3.1-Å resolution cryo-EM reconstruction of the dimeric TOM complex (TOM-pALDH). Tom subunits from each asymmetric unit are indicated by subscripts, A and B. Note that weak, low-resolution density features of the detergent micelle and the pALDH peptide are not shown. **c, d**, Atomic model of the TOM complex in ribbon representation. Two DDM detergent molecules between the Tom40 subunits are represented in sticks. Three α-helical segments (α1, α2, and α3) of Tom40 are indicated. Shown are a view from the cytosol (**a, c**) and a side view (**b, d**). Dotted lines (in **d**), outer membrane (OM) boundaries.

### Overall structure of the dimeric TOM complex

The near-atomic resolution density map of TOM-pALDH enabled us to build an accurate de novo model (Fig. 1c, d). A local resolution estimate indicates that a large portion of the complex, especially the Tom40 subunit, is at ∼3.0-Å resolution or better (Supplementary Fig. 2a). The map resolves not only individual β-strands of Tom40 but also almost all side chains (Supplementary Fig. 2c). Distal segments of Tom22 and small Tom subunits however remain poorly resolved likely due to intrinsic flexibility. Notably, our subunit assignment agrees with the previous assignment of the *N. crassa* structure^27^, which was largely based on crosslinking data^13^.

Each monomeric unit of the TOM complex contains a single copy of Tom40, Tom22, Tom5, Tom6, and Tom7 with each Tom40 forming a separate pore for polypeptide passage (Fig. 1). The new structure confirms that the Tom40 barrel consists of 19 β-strands (β1–19) arranged in an antiparallel fashion, except for β1 and β19, which are parallel. As noted previously^13,27^, an N-terminal segment (α2) spans the interior of the Tom40 barrel, exposing the N-terminus to the IMS. Both features, 19 β-strands and an N-terminal segment within the pore, closely resemble the structure of VDAC, despite low (∼15%) sequence identity^30^ (Supplementary Fig. 4). The two Tom40 subunits directly contact each other on the cytosolic side by hydrophobic side chains in β1-β19-β18 (Supplementary Fig. 5a–c). The interface is further wedged by the Tom22 helices (Fig. 1c, d, Supplementary Fig. 5d, e). Because the two Tom40 barrels are tilted away from each other by ∼40°, a gap exists in the Tom40-Tom40 interface which opens towards the IMS (Fig. 1d). In our structure, the gap is filled by two DDM detergent molecules as well as two Tom22 TMs (Fig. 1c, d, Supplementary Fig. 5c). In the native membrane, a phospholipid would occupy this gap in place of detergent with its headgroup phosphate positioned to interact with Arg330 of Tom40 (Supplementary Fig. 2c). The strong conservation of Arg330 suggests these lipids are important for dimer formation. The tilted arrangement does not seem to grossly bend the membrane based on the relatively flat micelle (Supplementary Fig. 6).

Tom22 contains an unusually long (∼45-amino-acid long) α-helix, the middle portion (roughly, positions 100–118) of which spans the membrane (Fig. 1d). The helix is longbow-shaped because of a kink formed by Pro112 (Fig. 2a), a residue that has been reported to be important for mitochondrial targeting of Tom22^31^ and stability of the TOM complex^13^. The helix extends at least 22 Å out from the membrane into the IMS, which may function as a binding site for presequences^32^ or the TIM complex^33^. On the opposite cytosolic side, the Tom22 helix becomes amphipathic, lying flat on the membrane surface (Supplementary Fig. 6). Preceding the helix, the cytosolic segment (positions 1–88) of Tom22 are invisible likely due to its flexibility. The function of this region has been suggested to be a docking site for Tom20 and Tom70 (ref. 34,35) and/or a presequence receptor^19,36^. The mechanism for the latter is unclear because the domain appears to be directed away from the Tom40 pores. The other three small Tom subunits (Tom5/6/7) are peripherally bound to Tom40 by interactions with different regions of Tom40 (Fig. 1). Although it appears as a low-resolution feature in our map, the N-terminal segment of Tom5 (∼12 amino acids) seems to be positioned above the Tom40 pore on the cytosolic side (Supplementary Fig. 6b), in agreement with its proposed role in recruiting precursor proteins to the pore^17^.

**Figure 2.**
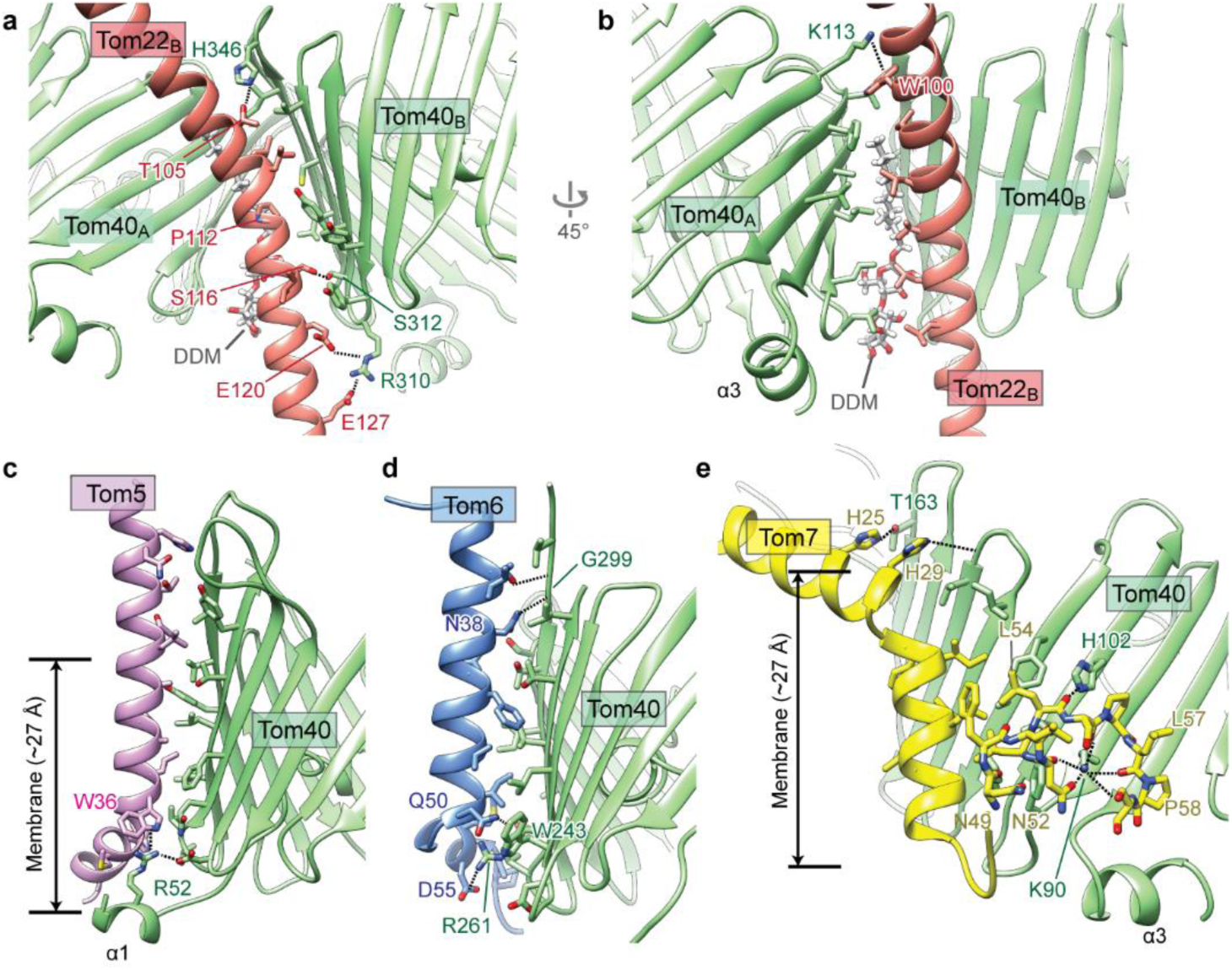
Inter-subunit contacts between Tom40 and α-helical Tom subunits. **a, b**, Interactions between Tom40 and Tom22 within the same monomeric unit (**a**; Tom40_B_–Tom22_B_) or between the different units (**b**; Tom40_A_–Tom22_B_). Amino acid side chains in the interfaces and the DDM detergent are displayed in stick representation. The polar interactions are indicated by black dotted lines. Shown are side views. **c**–**e**, As in **a, b**, but showing interactions of Tom40 with Tom5 (**c**), Tom6 (**d**), or Tom7 (**e**). Note that in **e**, N49–L54 of Tom7 is an α-helix.

### Interactions between β-barrel and α-helical membrane proteins

The TOM complex represents a rare example where a complex consists of both β-barrel and α-helical types of integral membrane proteins, and thus our structure offers a unique opportunity to examine interactions between the two types of membrane proteins. The structure shows that association between Tom40 and α-helical Tom subunits is mainly mediated by hydrophobic interactions in conjunction with high surface complementarity between transmembrane domains (Fig. 2, Supplementary Fig. 5d–h). In addition, several polar interactions were noticed near the membrane boundaries (Fig. 2). Conservation of these polar interactions across fungal species suggests that they may play an important role in increasing specificity and affinity of subunit interactions (Supplementary Table 1). Indeed, mutation of R261 or W243 of Tom40, which interacts with Tom6 in our structure, has been shown to decrease the stability of TOM similar to a Tom6 knockout^37,38^.

Our structure also reveals an interesting, unusual topology of Tom7, in which its C-terminal segment following the helical TM re-enters the membrane on the IMS leaflet (Fig. 2e). Part of the segment is unstructured and adopts a hook shape. An unstructured polypeptide in the lipid membrane is very rare because unpaired hydrogen-bond donors and acceptors of the peptide backbone would be energetically unfavorable. In the TOM complex, this issue seems to be overcome by hydrogen-bonding between backbone carbonyl oxygen atoms of Tom7 and lipid-facing side-chain nitrogen atoms of conserved Lys90 and His102 of Tom40. Although the exact function of Tom7 is unclear, its highly unique structure suggests a potential regulatory role in biogenesis and assembly of the TOM complex, as proposed previously^23^.

### Pore structure of Tom40

To gain insight into the protein translocation mechanism by TOM, we examined the translocation pathway in Tom40. While the Tom40 β-barrel has relatively large (∼30 Å by ∼25 Å) oval-shaped openings on both cytosolic and IMS sides, the pore is substantially constricted (∼19 Å by ∼13 Å) halfway across the membrane by the α2 segment (Fig. 1c). Still the pore would snugly fit one or perhaps two α-helices along the vertical translocation axis. Given the considerable contacts between α2 and the β-barrel interior and interactions of the preceding N-terminal amphipathic helix (α1) on IMS with Tom5 and the membrane (Figs. 1d, 2c), it seems unlikely that the α1-α2 segment becomes dislodged during protein translocation. It is also unlikely that the Tom40 barrel opens laterally towards the lipid phase as proposed for BamA and Sam50, which mediate membrane insertion of β-barrel proteins^39-41^. The only separable β-stand pair, β1-β19, is sealed by ∼10 hydrogen bonds, which would be energetically costly to split (not shown). Together with our observation that binding of a presequence did not cause any noticeable structural changes, these suggest that Tom40 is a static pore for polypeptide passage.

To understand how Tom40 may interact with translocating polypeptides, we evaluated surface properties of its pore (Fig. 3). Surface electrostatic analysis indicates that the Tom40 pore has an overall negative potential, mainly attributed to several acidic patches (APs 1–3) on the pore lining (Fig. 3a–d and Supplementary Fig. 7). A similar negative electrostatic potential is anticipated for Tom40 from other fungal species based on homology modelling (Supplementary Fig. 8). This explains why Tom40 is selective for cations when ion conduction was measured by electrophysiology^11,42^. The negative electrostatic potential likely promotes protein translocation by attracting positively-charged amino acids in polypeptides, such as inner membrane proteins and presequences of matrix-targeted preproteins, both of which are often basic^43^. Interestingly, the potential seems more negative towards the IMS side (Supplementary Fig. 8e), which may promote polypeptide movement towards IMS. The pore-lining surfaces also contain hydrophobic patches (particularly, patches labelled HP2 and HP3; Fig. 3e–h). These patches may interact with hydrophobic regions of precursor proteins to facilitate translocation.

**Figure 3.**
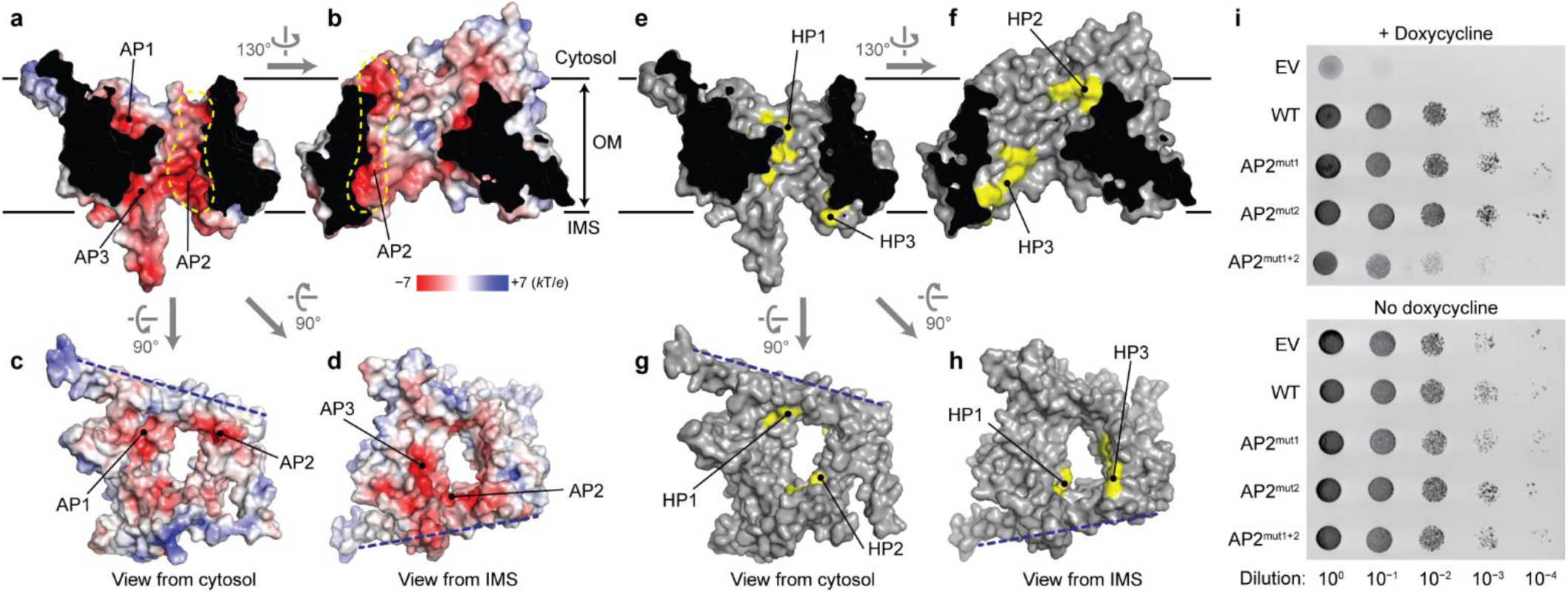
Pore architecture of Tom40. **a**–**d**, Surface electrostatics of the TOM complex shown as a heat map on a solvent-accessible surface representation. For simplicity, only one monomeric unit is shown (the dimer interface indicated by a blue dashed line). Shown are cutaway side views (**a, b**) and views from the cytosol (**c**) and IMS (**d**). Acidic patches are referred to as AP1, AP2 (also outlined by yellow dash line), and AP3. OM, outer membrane. **e**–**h**, As in **a**–**d**, but showing hydrophobic patches (HPs) in yellow. **i**, Yeast cells expressing wild-type (WT) or indicated mutant Tom40 from a plasmid were serially diluted and spotted on SC(−Ura/−Leu) plates containing or lacking doxycycline. In these strains, the presence of doxycycline represses expression of chromosomal Tom40. EV, empty vector. AP2^mut1^=D132N/D134N/E329N; AP2^mut2^=D87N/E360N; AP2^mut1+2^= D87N/D132N/D134N/E329N/E360N.

To test the functional importance of these patches, we examined cell growth defects accompanied by their mutations on the basis that Tom40’s protein translocation function is essential for cell viability. When we mutated the conserved acidic patch AP2 by replacing five Glu and Asp with Asn (mut1+2), substantial growth retardation was observed, whereas partial mutations (mut1 and mut2) did not impair growth (Fig. 3i). This suggests that the defect is likely due to loss of the negative potential in AP2. We also observed mild growth inhibition when we simultaneously mutated multiple hydrophobic amino acids in HP2 or HP3 (Supplementary Fig 7h). Relatively mild impairment might be due to functional redundancy of multiple patches in the pore.

### Binding of a presequence to the Tom40 pore

To understand the mechanism of presequence recognition by Tom40, we analyzed the pALDH peptide density in our cryo-EM map. To accurately define the density feature corresponding to pALDH peptide, we generated a difference map by subtracting intensity values of the apo TOM map from those of the pALDH-TOM map (Fig. 4a, b, Supplementary Fig. 9). The pALDH density is rather weak perhaps because of its heterogeneous position within the pore as well as low occupancy of the peptide, and therefore we imposed a lowpass filter to suppress noise. This revealed an elongated feature in the pore cavity of each Tom40, which we modelled as an 18-residue-long polyalanine α-helix (Fig. 4a, b).

**Figure 4.**
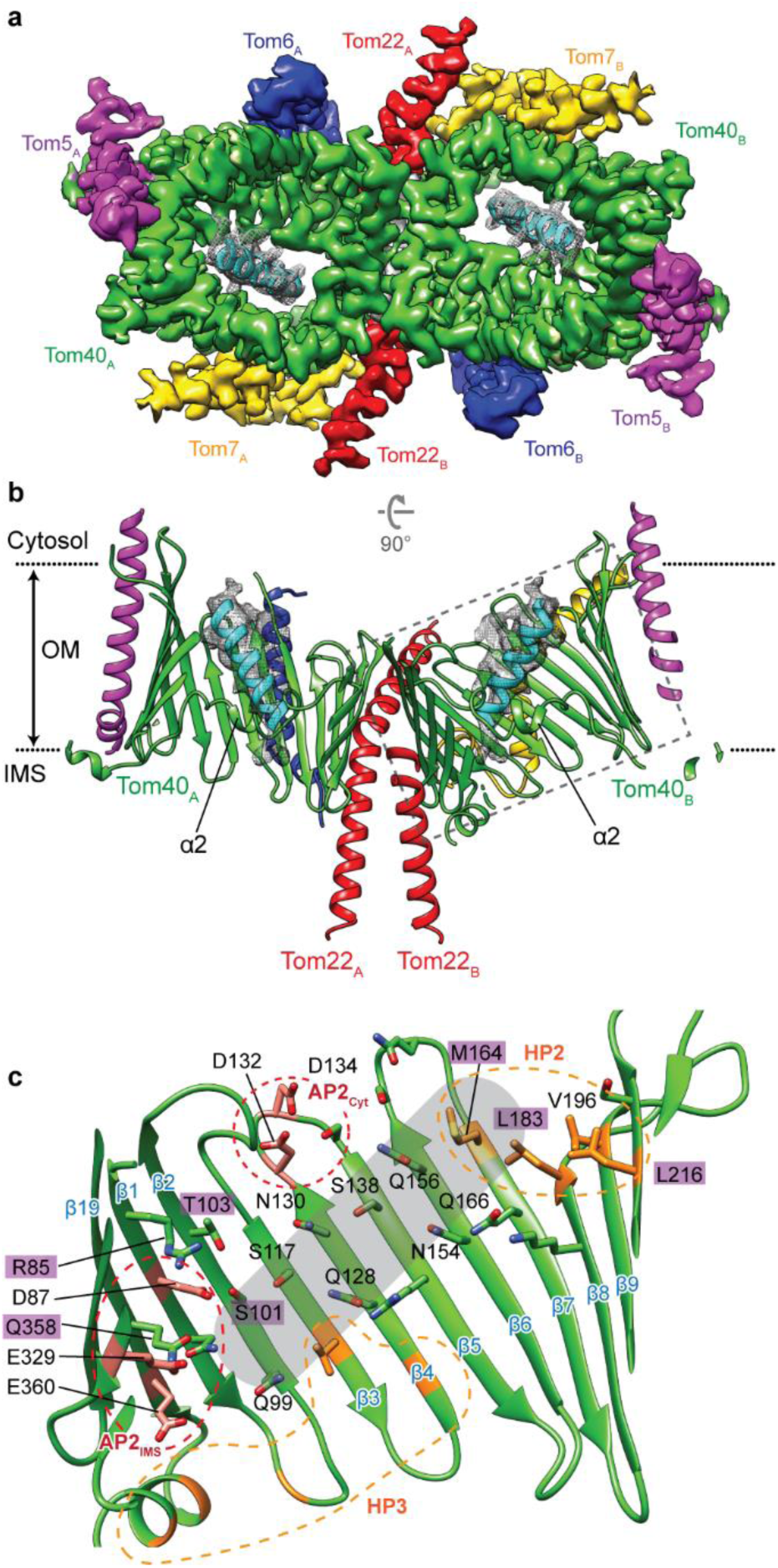
Presequence peptide binding to the TOM40 pore. **a**, Composite map (cytosolic view) showing the density for the pALDH peptide (grey mesh; difference map between the TOM-pALDH and apo TOM maps) overlaid on the TOM complex map in Fig. 1**a**. A model of pALDH is represented as a cyan ribbon. **b**, As in **a**, but showing a side view with the TOM complex in a ribbon model instead of the density map (as in Fig. 1**d**). The grey dashed line indicates the area shown in **c**. OM, outer membrane. **c**, Presequence-interacting surface of Tom40 (cutaway side view). Amino acid side chains near the bound pALDH peptide are shown in stick representation. A surface in immediate vicinity to the pALDH helix is shaded in grey. Acidic and hydrophobic residues are in salmon and orange, respectively. Residues that were previously shown to crosslink translocating substrates^13^ are highlighted in purple. For clarity, α2 is not shown. AP2, acidic patch 2. HP2 and HP3, hydrophobic patch 2 and 3, respectively (see Fig. 3).

The presequence is tilted ∼45° from the barrel axis and lies on the relatively flat surface formed by β2 to β7 with its direction nearly perpendicular to the β strands (Fig. 4c). The surface is relatively neutral and hydrophilic, lined with side chains of Gln, Asn, and Ser. On the opposite side, the presequence also seems to make a contact with α2 (Q78 and Y79; not shown). In addition, each end of the presequence helix appears to interact with acidic (AP2) and hydrophobic (HP2) patches of Tom40. However, due to insufficient side-chain features, we could not register specific amino acids into the pALDH density, and thus the exact nature of interactions between Tom40 and the presequence remains unclear. We speculate that the hydrophilic β2–7 surface might preferentially interact with the polar or positively charged amino acids of the presequence, and AP2 might interact with the N-terminus of the presequence (assuming physiological ‘head-first’ insertion) by a helical dipole moment.

While the presequence density spans along the pore, its position is closer to the cytosolic entry than to IMS. Therefore, our structure likely represents an early stage of preprotein engagement with Tom40. Previous planar lipid bilayer electrophysiology studies have shown that a presequence peptide binds to Tom40 with higher affinity when added from the cytosolic side than when added from the IMS side^11^. The pALDH position shifted towards the cytosol in our structure may explain this observation. Furthermore, many amino acids in Tom40’s presequence binding surface identified in our structure (e.g., R85, S101, T103, M164, L183, L216, and Q385; Fig. 4c) have previously been shown to crosslink to stalled translocation substrates^13^. This may suggest that various substrate polypeptides preferably interact with this surface.

### Assessment of oligomeric structure of TOM

A longstanding puzzle has been the oligomeric state of the TOM complex. It has been generally believed that the mature or holo TOM complex is a trimer. However, our structure suggests that the dimer is a stable configuration, and more importantly, translocation-competent. Given extensive inter-subunit contacts at the dimer interface, it is seemingly improbable that the dimer rearranges into a homotrimer. One possibility would be that binding of the Tom20 and Tom70 subunits induces trimerization of the TOM complex^28,29^, but we suggest that such a major subunit rearrangement is unlikely considering their low-affinity association^6,9^. Instead, we considered a possibility that detergent used in protein purification may affect the oligomeric state of the complex. To test this, we performed experiments under different detergent conditions, carefully monitoring SEC elution profiles to evaluate the size of the complex (Fig. 5 a– c, Supplementary Fig. 10).

**Figure 5.**
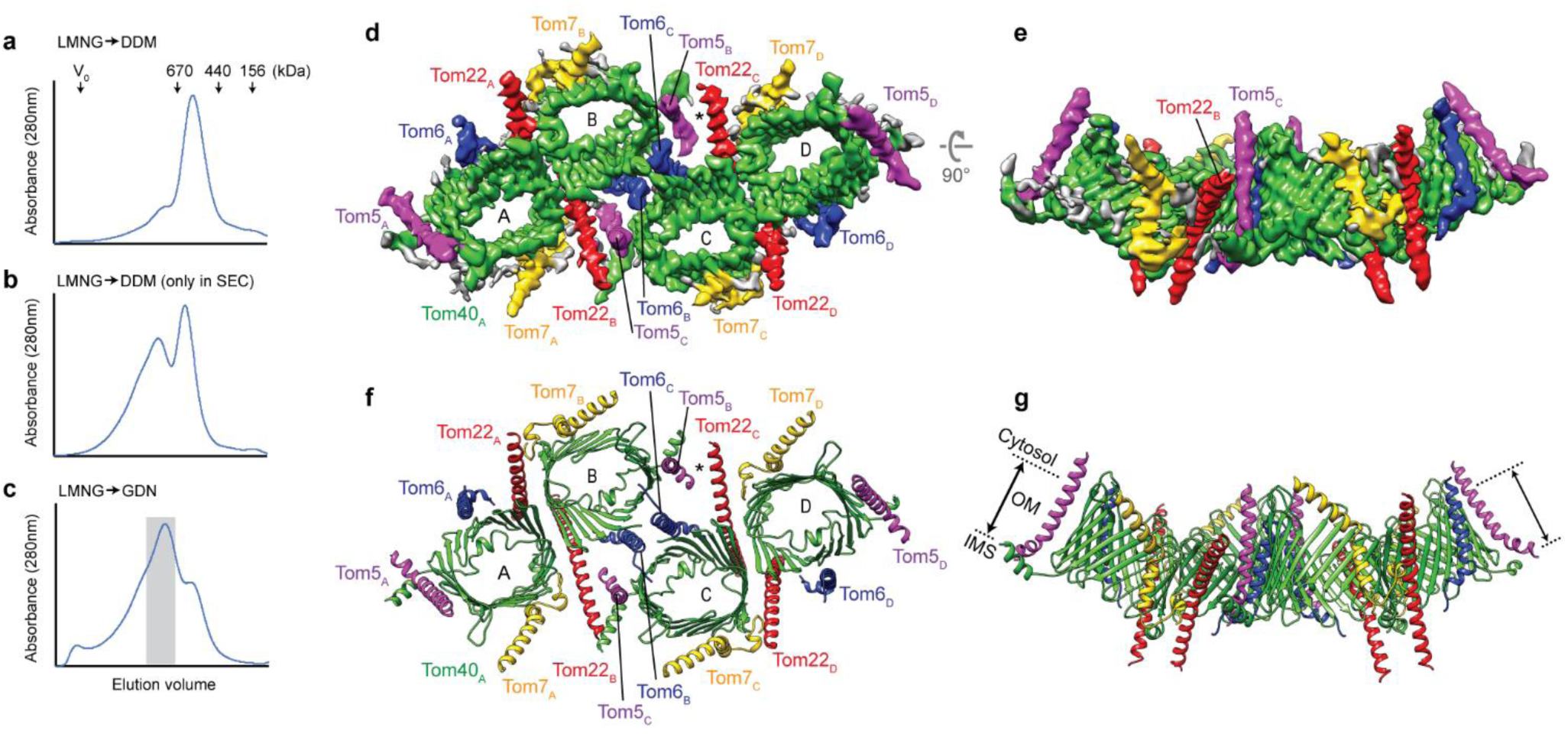
Cryo-EM structure of the tetrameric TOM complex. **a**–**c**, SEC elution profiles of the TOM complex in different detergent conditions (for details, see Supplementary Fig. 9). V_0_, void volume. In **c**, fractions in grey were used for the cryo-EM structure shown in **d**–**g. d**–**g**, Cryo-EM reconstruction (**d, e**) and atomic model (**f, g**) of the tetrameric TOM complex. Four monomeric units are indicated by A, B, C, and D. Shown are a view from the cytosol (**d, f**) and a side view (**e, g**). Asterisk, gap between Tom5_B_ and Tom22_C_. Dotted lines (in **g**), outer membrane (OM) boundaries.

Surprisingly, these experiments indicate that the dimeric TOM complex is indeed a product of dissociation of larger complexes. While exchange of LMNG into DDM during affinity purification resulted in almost exclusively dimers that migrated as an ∼500-kDa species (Fig. 5a), delayed exchange into DDM at the last SEC step produced an additional peak appearing at a higher molecular size (∼1 MDa) (Fig. 5b). When DDM was substituted by glyco-diosgenin (GDN), a digitonin-like detergent that is generally considered to be more gentle than DDM, the complex eluted mostly in the 1-MDa peak (Fig. 5c). The sample also seemed to contain even larger species as some TOM proteins eluted earlier. Importantly, SDS-PAGE analysis of peak fractions showed no changes in subunit composition (Supplementary Fig. 10f), indicating that the two peaks simply differ in their oligomeric states. Because many previous studies evaluating the TOM complex assembly have used blue native PAGE (BN-PAGE) analysis^6,9,10,19,22,34^, we also subjected crude extracts prepared with different detergents to both BN-PAGE and SEC analyses for comparison (Supplementary Fig. 11). These experiments suggest that the previously reported 400-kDa band in BN-PAGE analysis corresponds to the dimeric TOM complex. Unlike SEC analysis, however, BN-PAGE did not show prominent higher-oligomer species. It is possible that harsh conditions of BN-PAGE led to dissociation of higher oligomers into dimers^7^.

### Cryo-EM structure of the tetrameric TOM complex

To elucidate the structure of high-order TOM oligomers, we analyzed 1-MDa peak fractions by cryo-EM (Fig. 5, Supplementary Fig. 12). As expected, particles on micrographs were much larger than those seen with the dimer samples (Supplementary Fig. 12b). 2D and 3D classifications of particle images showed a striking tetrameric arrangement of the pores (Supplementary Fig. 12a, c). We also noticed that micrographs often showed particles larger than the dimensions of the tetramer, indicating that the sample included even larger oligomers (Supplementary Fig. 12g). Nonetheless, tetramers were the predominant species, and ∼80% of particles could be used for the final reconstruction of the tetrameric TOM complex at 4.1-Å resolution (Fig. 5d, e). Interestingly a minor 3D class showed three pores (Supplementary Fig. 12a; Class 3), reminiscent of trimers seen in low-resolution EM studies^5,28,29^. This ‘trimer’ class appears to be derived from the tetramer by dissociation of one monomeric unit.

The structure reveals that the tetramer is essentially a dimer of two dimeric TOM complexes (referred to as A/B and C/D), which are arranged in a staggered parallel fashion with units B and C being associated to each other (Fig. 5, Supplementary Fig. 13). There are only a few structural differences between the dimeric complex and dimers in the tetrameric complex as two copies of atomic models for the dimer could be fitted into the EM map essentially as rigid bodies. The contacts between units B and C are made mainly by the two Tom6 (Tom6_B_ and Tom6_C_) subunits that are sandwiched by the two Tom40 (Tom40_B_ and Tom40_C_) subunits (Fig. 5c, Supplementary Fig. 13e, f). Additionally, the N-terminal segment (residues 1–24) of Tom6 appears to be directed to the neighboring Tom40’s barrel interior next to β11 and near HP2 (Supplementary Fig. 14). There is also a minor contact between Tom22_B_ and Tom5_C_ where their TMs cross each other on the cytosolic membrane boundary. Interestingly the tetrameric interface is not completely symmetric, because a gap exists between Tom22_C_ and Tom5_B_ unlike the Tom22_B_–Tom5_C_ contact (Fig. 5f, Supplementary Fig. 13 a–c). Furthermore, there is a considerable gap (∼7 Å in width) along the dimer-dimer interface at the IMS leaflet of the membrane (Supplementary Fig. 13g, i). In the cryo-EM map, the gaps are filled by weak density features, which should be detergent and/or lipid molecules (Supplementary Fig. 13h and data not shown). The relatively loose interface explains why tetramers easily dissociate into dimers by excess detergent.

The observed subunit arrangement suggests that the complex can further expand into larger oligomers in the membrane, which is further supported by observations of a broad peak width in SEC analysis and larger particles in EM micrographs (Supplementary Fig. 10c–e and 12g). Because of the crevice opened to the IMS along the dimer-dimer interface, oligomerization might create a curvature (concave to the cytosolic side) of ∼30° per added dimer (Fig. 5d). However, it is possible that in the native membrane, the gap is closed such that the complex lies relatively flat in the membrane. Looking from IMS, protein surfaces in the interface are roughly complementary between the two TOM dimers to accommodate such a closure (Supplementary Fig. 13i).

## Discussion

Our study offers new mechanistic insights into how Tom40 initiates translocation of precursor proteins. While precursor polypeptides are first recognized by the cytosolic domains of Tom20, Tom22, and Tom70, they need to be threaded into the pore of Tom40. Because there is no external energy input (i.e., ATP or membrane potential) involved, the process must be driven solely by affinity of precursor proteins towards the pore interior. Our structure shows that, in case of at least one matrix-targeted protein, its N-terminal presequence helix inserts into the Tom40 pore mainly by electrostatic and polar interactions. This binding mode may provide an additional ‘filter’ for increased targeting specificity because initial recognition of presequences by Tom20 is mediated by hydrophobic interactions^18^. Our findings also suggest an explanation for why mitochondrial targeting presequences are considerably less hydrophobic than the endoplasmic reticulum (ER)-targeting signal sequences of secretory proteins. While recognition of signal sequences by the Sec61 ER protein-translocation channel involves their partitioning into the hydrophobic lipid phase of the membrane^44,45^, mitochondrial presequences remain in largely hydrophilic environments throughout their insertion into and translocation across Tom40.

A highly unexpected finding was that the TOM complex forms a tetramer and larger oligomers instead of the trimer as previously thought. Our study also revealed that formation of the TOM tetramer is mediated by Tom6. This new structural insight about the involvement of Tom6 coincides well with its proposed function in stabilizing the TOM complex in vivo^46^. Consistent with the role we are now able to ascribe to Tom6, it has been shown previously that phosphorylation of Tom6’s N-terminal tail (Ser16) increases the steady-state level of Tom6 and the TOM complex as well as overall mitochondrial protein import^46^. Oligomerization may improve import efficiency by clustering of Tom40 pores, particularly when protein import occurs cotranslationally, during which many precursor molecules would be produced locally by polysomes^47,48^. In addition, the orientation of Tom6’s N-terminal segment toward the Tom40 pore may suggest a ‘hand-off’ function that facilitates interaction of the Tom40 pore with precursor proteins, thereby increasing translocation efficiency. Finally, our work provides a framework for further investigations to understand the structure, dynamics, and functions of the high-order TOM complex assemblies we have discovered.

## Acknowledgments

We thank D. Toso for help with electron microscope operation and J. Thorner for yeast strains and antibodies. We thank J. Thorner, J. Hurley, S. Brohawn, and S. Itskanov for critical reading of manuscript. This work was funded by U.C. Berkeley and the Vallee foundation (E.P.).

## Author information

### Contributions

E.P. conceived the project. K.T. and E.P. performed experiments. E.P. built the atomic models. K.T. and E.P. interpreted results and wrote the manuscript. E.P. supervised the project.

### Competing interests

The authors declare no competing interests.

### Data availability

The cryo-EM density maps and atomic model will be available through EM DataBank and Protein Data Bank, respectively.

## Methods

### Constructions of plasmid and yeast strains

To generate an *S. cerevisiae* strain overexpressing the TOM complex components from an inducible GAL1 promoter, we used the Yeast Tool Kit (YTK) and Golden Gate assembly^49^. We first amplified coding sequences (CDS) for Tom40, Tom22, Tom20, Tom 7, Tom6, and Tom5 by PCR using genomic DNA of *S. cerevisiae* BY4741 as a template and cloned them individually into the pYTK1 entry plasmid. To enable affinity purification of the Tom complex, a Strep-tag (GGWSHPQFEK) and a His-tag (GGHHHHHHHH) were introduced before the stop codons of Tom40 and Tom22, respectively. The cloned Tom subunits were combined with YTK parts to generate individual expression cassettes, each containing the *GAL1* promoter (YTK30), CDS of a Tom subunit, and the *ENO1* terminator (YTK61). We then assembled the six expression cassettes into a single multigene plasmid concatenating them in the order of Tom40-Tom22-Tom20-Tom7-Tom6-Tom5. The plasmid also contained a nourseothricin resistance marker (YTK78) for selection and URA3 homology arms (YTK92 and YTK86) for chromosomal integration. The resulting assembly was introduced to the YMLT62 yeast strain (a gift from J. Thorner) by a standard lithium acetate transformation method after linearizing the plasmid with the NotI endonuclease. The colonies were selected on a YPD agar plate containing 100 μg/mL nourseothricin, and chromosomal integration was confirmed by PCR. The YMLT62 strain (BY4741 leu2::pACT1-GEV::HIS3MX) contains the chimeric transcriptional activator Gal4dbd.ER.VP16 (GEV; ref. 50) integrated to the *LEU2* locus, which induces the transcription by the *GAL1* promoter upon addition of β-estradiol to the growth medium.

To generate plasmids for yeast Tom40 complementation tests, we first amplified by PCR the endogenous Tom40 gene region (of BY4741) containing the 329-bp upstream segment of the start codon and the 381-bp downstream segment of the stop codon. This fragment was then inserted into a home-made yeast CEN/ARS plasmid constructed with YTK (used parts: pYTK84, pYTK8, pYTK47, pYTK73, pYTK75, and pYTK81). The plasmid contains a LEU marker for selection. For immunodectection, we attached a Strep-tag to the C-terminus of Tom40 by PCR. Indicated mutations were introduced by PCR.

### Purification of the TOM complex

Yeast cells were grown in YPEG medium (1% yeast extract, 2% peptone, 2% ethanol and 3% glycerol) in shaker flasks at 30°C. Upon reaching an optical density (OD600) of ∼1.4–2, cells were induced with 50 nM β-estradiol. After 9–10 h of induction, cells were harvested by centrifugation at 5,000 rpm. Cell pellets were flash-frozen in liquid nitrogen and stored in −80°C until use. The TOM complex was purified by tandem affinity purification using His-and Strep-tags as summarized in Supplementary Fig. 10a. Cells were first lysed by cryo-milling (SPEX SamplePrep) at the liquid nitrogen temperature and resuspended in buffer (3 times cell pellet volume) containing 50 mM Tris-HCl pH 8, 200 mM NaCl, 10% glycerol, 20 mM imidazole, and protease inhibitors (5 µg/ml aprotinin, 5 µg/ml leupeptin, 1 µg/ml pepstatin A, and 1 mM PMSF). Then, one cell pellet volume of 5% lauryl maltose neopentyl glycol (LMNG; Anatrace) and 1% cholesteryl hemisuccinate (CHS; Anatrace) was added to solubilize membranes. After 3-h incubation at 4°C, the lysate was clarified by ultracentrifugation (Beckman Coulter rotor Type 45 Ti) at 125,000g for 1 h. The lysate was incubated by gentle rotation with HisPur cobalt resin (Life technologies) for 3 h at 4°C. The beads were then packed in a gravity column and washed with approximately 10 column volumes (CV) of buffer containing 50 mM Tris-HCl pH 8, 200 mM NaCl, 0.02% LMNG-CHS, 20 mM Imidazole and 10% glycerol. Resin was further washed with an additional 10 CV of buffer containing 40 mM imidazole and eluted with approximately 6 CV of buffer containing 180 mM imidazole. The eluate was then mixed with Strep-Tactin Sepharose (IBA Lifesciences) for ∼14 h at 4°C. The beads were packed in a gravity column and washed with approximately 10 CV of buffer containing 20 mM Tris-HCl pH 7.5, 100 mM NaCl, 0.03% dodecyl-β-maltoside (Anatrace), 0.006% CHS, and 1 mM dithiothreitol (DTT). In the case of purification of the tetrameric TOM complex, 0.02% glyco-diosgenin (GDN; Anatrace) was used instead of DDM/CHS. The TOM complex was eluted with buffer containing 3 mM D-desthiobiotin, and concentrated using AmiconUltra (100kDa cut-off, Millipore). The complex was further purified by SEC using a Superose 6 Increase column (GE Lifesciences) equilibrated with 20 mM Tris-HCl pH 7.5, 100 mM NaCl, 1 mM DTT, and 0.03% DDM/CHS (for the dimeric TOM complex) or 0.02% GDN (for the tetrameric TOM complex). Peak fractions were pooled, concentrated to ∼3.5–5 mg/mL using AmiconUltra (100kDa cut-off), and used to prepare cryo-EM grids. For experiments described in Supplementary Fig. 10 b–f, essentially the same procedure was employed but with modified detergent conditions as indicated.

### Cryo-EM specimen preparation and data acquisition

Immediately before preparing cryo-EM grids, 3mM fluorinated Fos-Choline-8 (FFC8; Anatrace) was added to the purified TOM sample. In the case of the presequence-bound Tom complex, before adding FFC8, a chemically synthesized pALDH peptide was added to the purified TOM complex at 10-fold molar excess with respect to Tom40 (incubated at 4°C for 30 min). We note that the addition of 3 mM FFC8 did not cause any changes in the SEC profiles of either the dimeric or tetrameric TOM complex even after a prolonged (∼6 h) incubation. To prepare cryo-EM grids, ∼3 μL of the sample was applied to a glow-discharged Quantifoil holey carbon grid (R 1.2/1.3 Au, 400 mesh; Quantifoil). Glow discharge was carried out for 20 s in 75% argon and 25% oxygen using a Gatan Solarus plasma cleaner or in air using a PELCO easiGlow glow discharge cleaner. The grid was blotted with Whatman No. 1 filter papers for 3 s at 4°C and 100% humidity and plunge-frozen in liquid-nitrogen-cooled liquid ethane using Vitrobot Mark IV (FEI).

A summary of image acquisition parameters is shown in Supplementary Table 2. For the dimeric TOM-pALDH complex and the apo tetrameric TOM complex, the datasets were collected on a Titan Krios electron microscope (FEI) equipped with a K2 Summit direct electron detector (Gatan) and a GIF Quantum image filter (Gatan). The microscope was operated at an acceleration voltage of 300 kV. Does-fractionated images were collected in the super-resolution mode with a physical pixel size of 1.15 Å and a GIF slit width of 20 eV using SerialEM software^51^. The dose rate was 1.22 electrons/Å^2^/frame with the frame rate of 0.2 s. For the TOM-pALDH complex, the total accumulated dose was 61 electrons/Å^2^ (50 frames), and for the tetrameric TOM complex, it was 48.8 electrons/Å^2^ (40 frames). The datasets for the apo dimeric TOM complex were collected similarly but using a Talos Arctica electron microscope (FEI) operated at an acceleration voltage of 200 kV and equipped with a K2 detector (without GIF). The images were recorded at physical pixel size of 1.16 Å with a dose rate of 1.25 electrons/Å^2^/frame (0.2 s/frame) and the total accumulated dose of 50 electrons/Å^2^ (40 frames).

### Single-particle image analysis of the TOM-pALDH complex

A summary of the single-particle analysis procedure is described in Supplementary Fig. 1c. Briefly, RELION3 (ref. 52) was used for preprocessing of movies, particle picking, and Bayesian particle polishing, and then cryoSPARC v2 (ref. 53) was used for ab-initio reconstruction, 3D classification, and the final 3D reconstruction. First, the movies were imported to RELION3 and corrected for motion using MotionCor2 with 5-by-5 tiling (ref. 54). During this step, micrographs were 2x-pixel-binned (resulting in a pixel size of 1.15 Å). Micrographs that were not suitable for image analysis (e.g., micrographs containing crystalline ice or displaying a large drift) were removed by manual inspection. Defocus parameters were estimated using CTFFIND4 (ref. 55). Template-based automatic particle picking was performed in RELION3 (460,148 particles from 1,587 movies). The particle templates were generated by 2D classification from Laplacian auto-picking on a subset of the data. The particles were extracted from micrographs with the box size of 256 pixels. Reference-free 2D classification (Supplementary Fig. 1d) was performed to remove empty detergent micelles and obvious non-protein particle artefacts, resulting in 290,793 particles. The initial 3D model was generated by cryoSPARC (ab initio reconstruction). The first 3D refinement was carried out by RELION3 using a lowpass-filtered initial model and 290,793 particle images, yielding a 3.8-Å resolution reconstruction. The particle images were subjected to one round of CTF refinement and Bayesian particle polishing in RELION3. These particles were subjected to second 3D refinement, which yielded 3.6-Å resolution reconstruction. Then, another round of CTF refinement and particle polishing was performed. The resulting polished particles were imported to cryoSPARC v2 for the subsequent process as described below.

The imported particles were subjected to 2D classification in cryoSPARC to further discard artefacts and low-quality particles. The resulting 243,227 particles were used to generate four ab initio 3D reconstructions, followed by heterogeneous refinement (3D classification). 179,232 (74%) particles converged to one class (Class 3; Supplementary Fig. 1c) leading to a high-resolution reconstruction of the dimeric TOM complex, whereas two low-resolution classes (Classes 1 and 2) appeared to have only a single pore, likely corresponding to dissociated monomers. After a second round of 3D classification to further remove low-quality particles, 160,577 from Class 3 were refined by non-uniform refinement with C2 symmetry imposed, yielding the final map at 3.06-Å resolution (based on gold-standard Fourier shell correlation (FSC) and the 0.143 cut-off criterion; Supplementary Fig. 1e). Local resolution was estimated by cryoSPARC using default parameters (Supplementary Fig. 2a).

### Single-particle image analysis of the apo dimeric and tetrameric TOM complexes

Summaries of single-particle image analyses for the apo dimeric and tetrameric TOM complexes are shown in Supplementary Figs. 3a and 12a, respectively. Essentially, motion correction, defocus estimation, particle picking, and particle extraction were performed using Warp (ref. ^56^), and the remaining downstream refinement process was carried out using cryoSPARC v2. Movies were corrected for motion with 8-by-8 tiling and defocus parameters were estimated with 5-by-5 tiling. Original super-resolution micrographs were 2x-pixel-binned. Particles were automatically picked by Warp. Micrographs were manually inspected to remove unsuitable micrographs. Particle images were extracted with a box size of 256 (for dimeric TOM) or 400 (for tetrameric TOM) pixels from dose-weighted frames 1–36 (skipping the last 4 frames). Particle images were then imported to cryoSPARC and subjected to one round of reference-free 2D classification to remove empty micelles. Ab initio reconstruction was performed to generate three (for dimeric TOM) or four (for tetrameric TOM) initial 3D models, which were further refined by heterogeneous refinement. For both dimeric and tetrameric apo TOM complex, ∼80% particles images converged into one (Class 1 in dimeric TOM) or two classes (Classes 1 and 2 in tetrameric TOM) showing high-resolution features. These particle images were used for the final 3D reconstructions by non-uniform refinement in cryoSPARC, yielding maps at resolutions of 3.5 Å (dimeric TOM) and 4.1 Å (tetrameric TOM), respectively. In the case of dimeric TOM, C2 symmetry was imposed. For the tetrameric TOM complex, symmetry was not imposed because the complex was found not completely symmetric (imposition of C2 symmetry led to artificial distortion of some density features). Local resolution was also estimated by cryoSPARC using default parameters.

### Difference map

To identify the density attributed to the pALDH peptide, we generated a difference map (Supplementary Fig. 9). We first lowpass (4.0 Å)-filtered the dimeric apo TOM and TOM-pALDH maps by the *relion_image_handler* program. To further suppress high-frequency noise, we also applied B-factor blurring (50 Å^2^) to the maps. The resulting processed maps were aligned in UCSF Chimera (ref. 57) (using the *fitmap* and *vop resample* functions). The difference map was produced by the *vop subtract* function of Chimera. Because the maps were at different density scales (the TOM-pALDH map has roughly 2-fold higher voxel intensity values than the apo TOM map), before map subtraction, we scaled the apo map by a factor of 2.5 (a multiplicative factor of 2.0 to 2.5 produced similar good results).

### Atomic model building

A summary of model refinement and validation is shown in Supplementary Table 2. The atomic model for dimeric TOM was built de novo using Coot (ref. 58) and the summed map of the TOM-pALDH complex. In addition to proteins, we also modelled several hydrophobic tails of detergent or lipid (we used DDM as a model). The model was refined in real space using Phenix (ref. 59) and the summed map with the refinement resolution limit set to 3.1 Å. Different weights were tested using half maps to check whether the used Phenix refinement protocol shows overfitting to the map (Supplementary Fig. 2b; FSC_work_ vs FSC_free_). To this end, we chose a weight of 2, which did not separate FSC_work_ and FSC_free_. We also used restraints for secondary structure. The poly-alanine model for the pALDH peptide was fitted in the difference map using Coot and were not further refined. The following segments were not modeled because of poor or invisible density features: N–48, 277–294, and 374–387(C) of Tom40, N–85 and 136–152(C) of Tom22, N–12 and N– 26 and 48–50 (C) of Tom6, and N–10 of Tom7. The model for the apo dimeric TOM complex was generated by rigid-body docking of the TOM model (of TOM-pALDH) into the map and one round of refinement with Phenix against the apo dimeric TOM map.

To build a model for the tetrameric TOM complex, two dimer models were fit into the tetramer map using UCSF chimera. A few additional residues (α1 of Tom40, 81–89 of Tom22, and 25–26 of Tom6) were built using Coot because the tetramer map shows extra densities for these segments. In addition, we modelled 1,2-dimyristoyl-*rac*-glycero-3-phosphocholine (DMPC) into the density at the Tom40-Tom40 dimer interface (instead of DDM as in the dimeric TOM complex). The model was then refined against the tetramer map essentially the same as described for the dimeric TOM complex. Structural validation was done by MolProbity (ref. 60).

Protein electrostatics were calculated using PDB2PQR and the Adaptive Poisson-Boltzmann Solver (www.poissonboltmann.org; ref. 61) with monovalent mobile ions (0.1 M for both cation and anion) included in parameters. UCSF Chimera and PyMOL (Schrödinger) were used to prepare figures in the paper.

### Tom40 complementation assays

To deplete endogenous wild-type Tom40, we used a yeast strain (TH_7610; Dharmacon) from Yeast Tet-Promoters Hughes Collection, in which the original Tom40 promoter was replaced by a tetracycline (Tet)-promoter. The cells were transformed with a CEN plasmid constitutively expressing wild-type or mutant Tom40-Strep under the endogenous promoter and selected on agar plates of a synthetic complete medium containing 2% glucose and lacking uracil and leucine (SC(-Ura/-Leu)). After 3-day incubation at 30°C, colonies were isolated. Cells were grown in 3 mL of SC(-Ura/-Leu) at 30°C until OD600 reached ∼0.7–1.5, pelleted, and resuspended in fresh medium at OD600=1. After 10-fold serial dilution, 10 μL were spotted on SC(-Ura/-Leu) agar plates. Where indicated, 15 μg/mL doxycycline was included in the medium to repress endogenous Tom40 expression. Plates were incubated at 30°C for ∼2.5 days before imaging. To test expression of the Tom40 mutants in cells, an equal number (2 ODs) of cells were collected from cultures in SC(-Ura/-Leu) medium, and proteins were extracted by heating in NaOH/SDS buffer. The samples were analyzed by SDS-PAGE and immunoblotting with anti-Strep (Genscript; A01732) and anti-PGK1 (a gift from J. Thorner) antibodies. Standard enhanced chemiluminescence reagents and a Fujifilm LAS-3000 Imager were used for detection.

### Size-exclusion chromatography (SEC) and blue native PAGE (BN-PAGE) analysis of crude extracts

Yeast cells were grown in YPEG medium and induced by β-estradiol as previously stated. Cells from ∼10-ml induced culture were pelleted, washed in distilled water, frozen in liquid nitrogen and stored at −80°C until use. Pelleted cells (∼100 mg) were resuspended in 400 μL of lysis buffer containing 50mM Tris-HCl pH 7.5, 200mM NaCl, 1mM EDTA, 2 mM DTT, and protease inhibitors. Cells were lysed by beating with pre-chilled glass beads (2 cycles of 1.5-min beating and 1-min rest). Beads were removed, and the lysate was mixed with detergent (from a 5% stock solution) as indicated. After solubilizing membranes for 1 h at 4°C, samples were clarified for 1 h at 13,300 rpm and 4°C. 100 μl of the clarified sample was injected into Superose 6 Increase column equilibrated with 20 mM Tris-HCl pH 7.5, 150 mM NaCl, 1 mM EDTA, 1 mM DTT, and a low concentration of detergent used for lysis (i.e., 0.03% DDM/0.006% CHS, 0.02% LMNG/0.004% CHS, 0.02% GDN, or 0.08% digitonin). Fractions were collected and analyzed to SDS-PAGE and immunoblotting analyses. For immunoblotting, anti-Strep-tag (Genscript) and anti-His-tag (Life Technolgies; MA1-21315) monoclonal antibodies were used.

Samples for BN-PAGE were prepared essentially the same way but with a minor modification. The lysis buffer contained 50mM Tris pH 7.5, 50mM NaCl, 10% glycerol, 1mM DTT, and protease inhibitors. Detergent-solubilized lysates were clarified by ultracentrifugation for 30 min at 250,000g (Beckman TLA-100 rotor) and 4°C. Coomassie Blue G-250 (prepared as 5% stock in 0.5 M 6-aminohexanoic acid; 1/4 amount of added detergent by weight) was added to the lysate. BN-PAGE was performed using a 4–16% Novex Native PAGE gel (Life Technologies) according to manufacturer’s instructions.

