## Supplementary Figures and Tables for "Cryo-EM structure of the mitochondrial protein-import channel TOM complex from *Saccharomyces cerevisiae*"

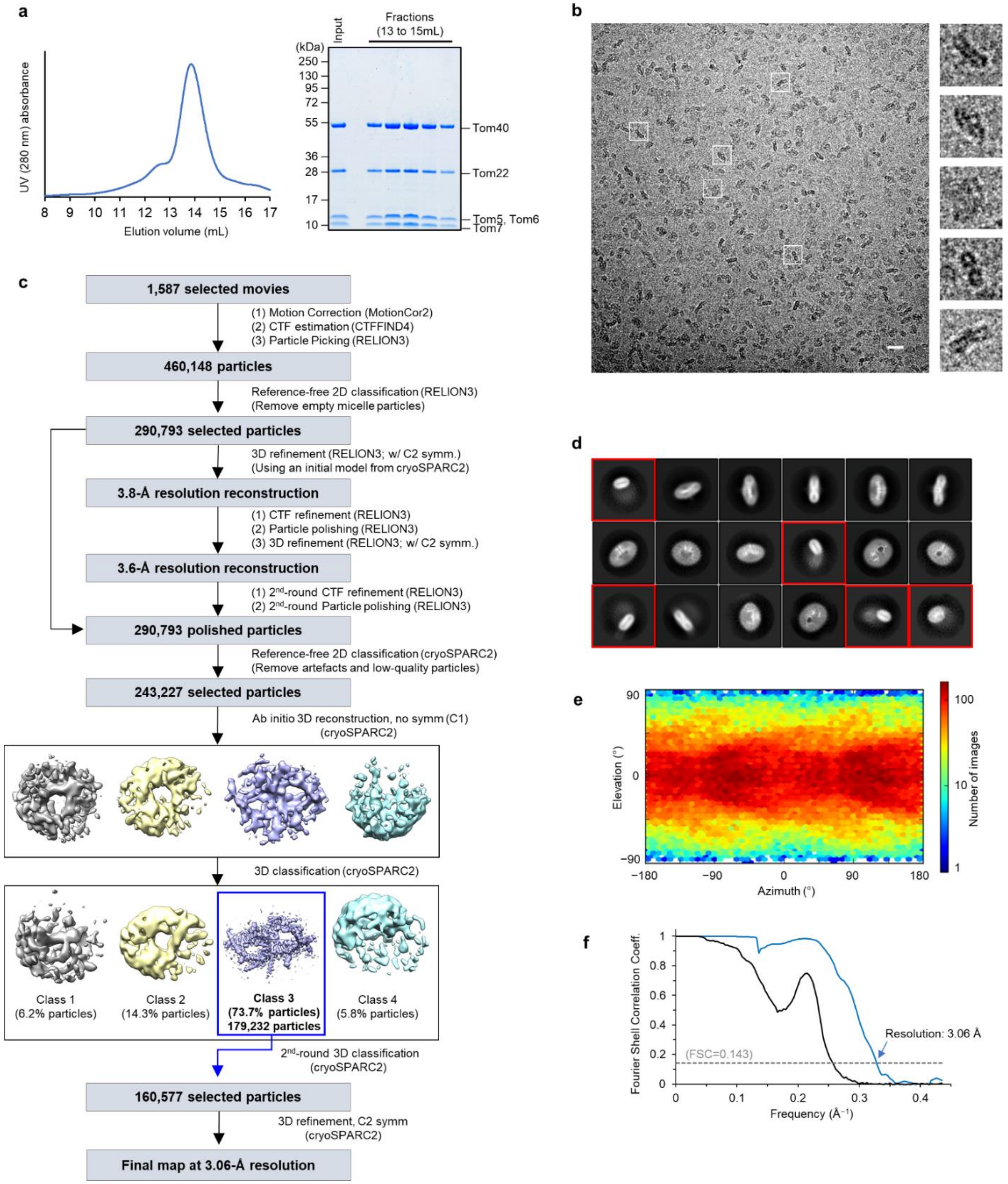


**Supplementary Figure 1. Single-particle cryo-EM analysis procedure for the presequence-bound dimeric core TOM (TOM-pALDH) complex.**

**a**, Size-exclusion chromatography (Superose 6) profile of affinity purified the yeast dimeric TOM complex. Left panel, absorbance at 280 nm. Right panel, Coomassie-stained SDS gel of peak fractions. **b**, A representative motion-corrected micrograph. Scale bar, 20 nm. Right panels show magnified images of selected particles outlined with white squares. The particle image size is 209 Å (width) by 209 Å (height). **c**, Summary of single-particle image analysis procedure. **d**, Representative class averages from 2D classification by RELION3 (the second step in **c**). The box dimensions are 297 Å (width) by 297 Å (height). Classes in red boxes are likely empty micelles and thus excluded in subsequent analysis. **e**, Heat map showing particle orientation distribution (produced in the final 3D reconstruction by cryoSPARC2). **f**, Fourier shell correlation (FSC) of two independently-refined half maps. Blue line, corrected masked FSC. Solid black line, unmasked FSC.

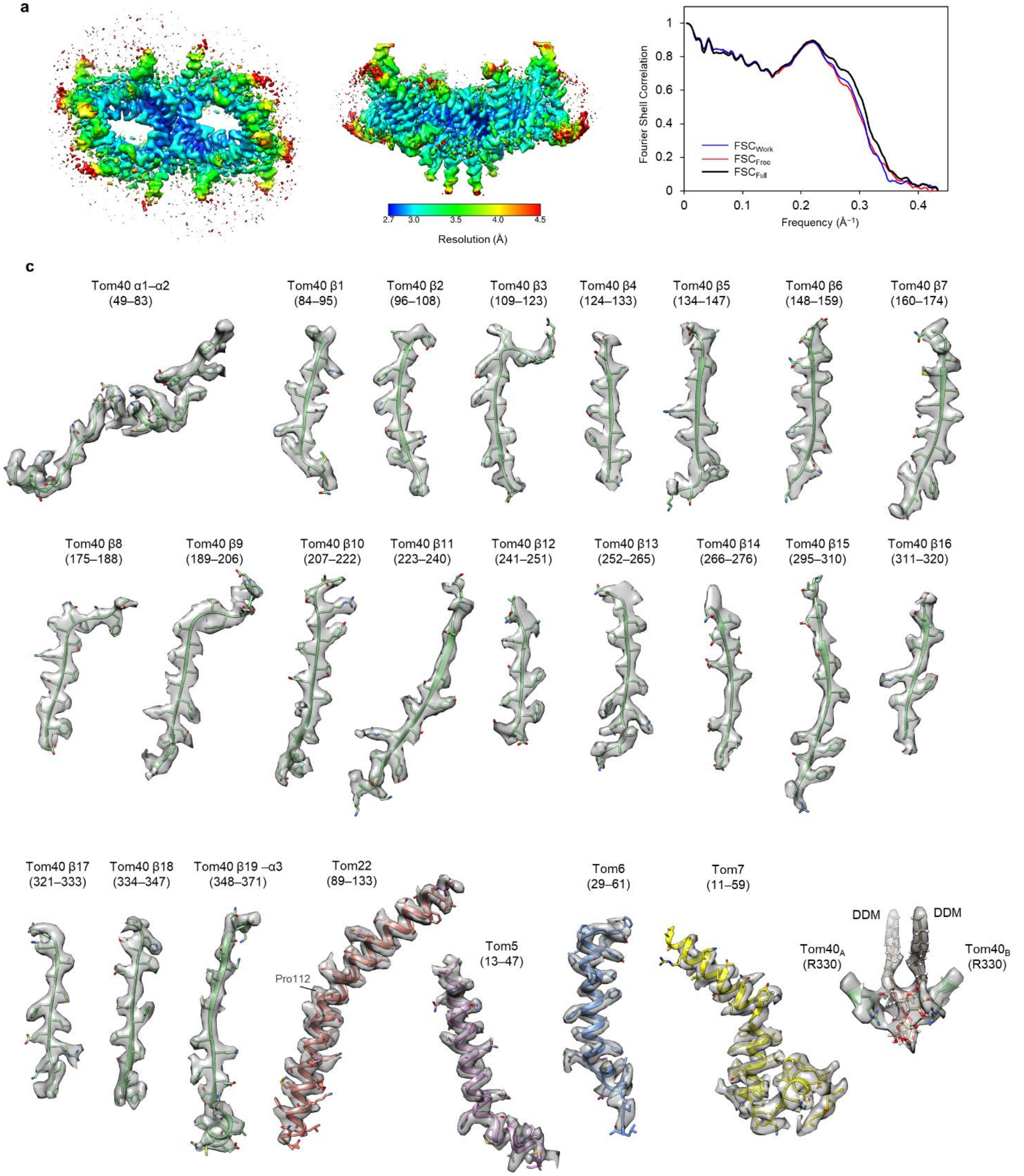


**Supplementary Figure 2. Cryo-EM map and atomic model quality of the TOM-pALDH complex.**
**a**, Local resolution is represented by a heat map on the density contour (unsharpened, summed map). **b**, FSC between the EM map and the atomic model. Blue curve, $FSC_{Work}$ (FSC between half map 1 and a model refined against half map 1). Red curve, $FSC_{Free}$ (FSC between half map 2 and the model refined against half map 1). Black curve, $FSC_{Full}$ (FSC between the combined map and the final atomic model refined against the combined map. All refinements were performed by Phenix with the same weight (2). **c**, Panels of the density map and the atomic model for indicated segments. Numbers in the brackets indicate ranges of amino acid residues shown.

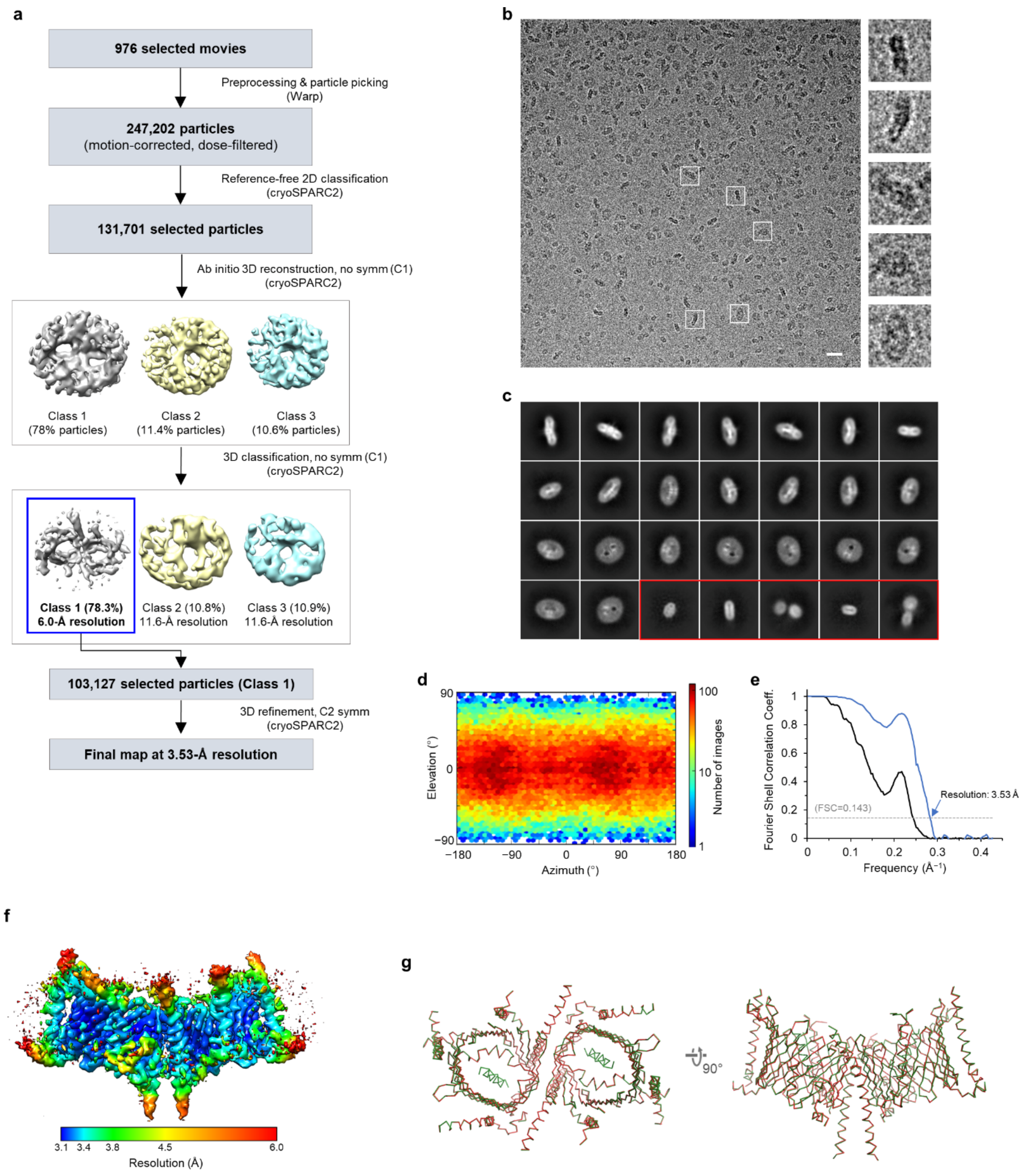


**Supplementary Figure 3. Single-particle cryo-EM analysis procedure for the apo dimeric core TOM complex.** **a**, Summary of single-particle image analysis procedure. **b**, A representative motion-corrected micrograph. Scale bar, 20 nm. Right panels show magnified images of representative particles outlined with white squares. The particle image dimensions are 209 Å (width) by 209 Å (height). **c**, Class averages from 2D classification (the second step in **a**). The box dimensions are 297 Å (width) by 297 Å (height). Classes in the red box likely represent empty micelles and thus excluded from further analysis. **d**, Particle orientation distribution. **e**, Fourier shell correlation (FSC) of two independently-refined half maps. Blue line, corrected masked FSC. Solid black line, unmasked FSC. **f**, Local resolution map (overlaid on the unsharpened summed map). **g**, Superimposition between the TOM-pALDH (green) and apo TOM (red) models. Left, view from cytosol; right, side view.

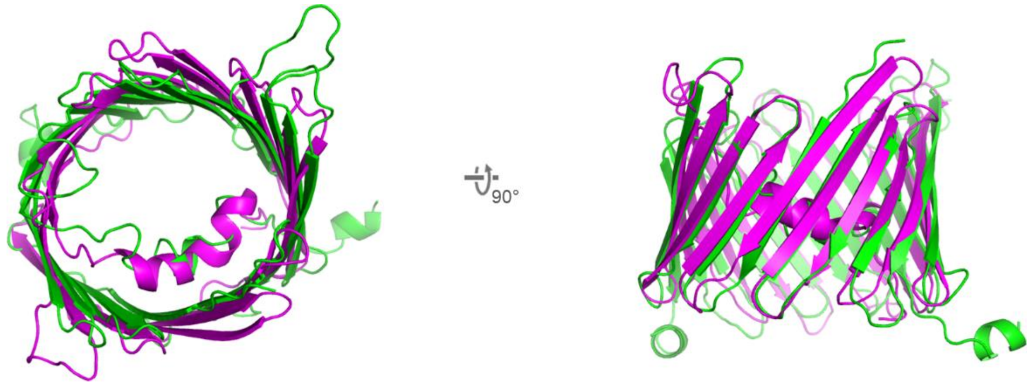


**Supplementary Figure 4. Structural comparison of Tom40 and VDAC.**
Structures of Tom40 (this study; green) and murine VDAC (PDB ID: 3EMN; magenta) are superimposed. Left, view from cytosol. Right, side view.

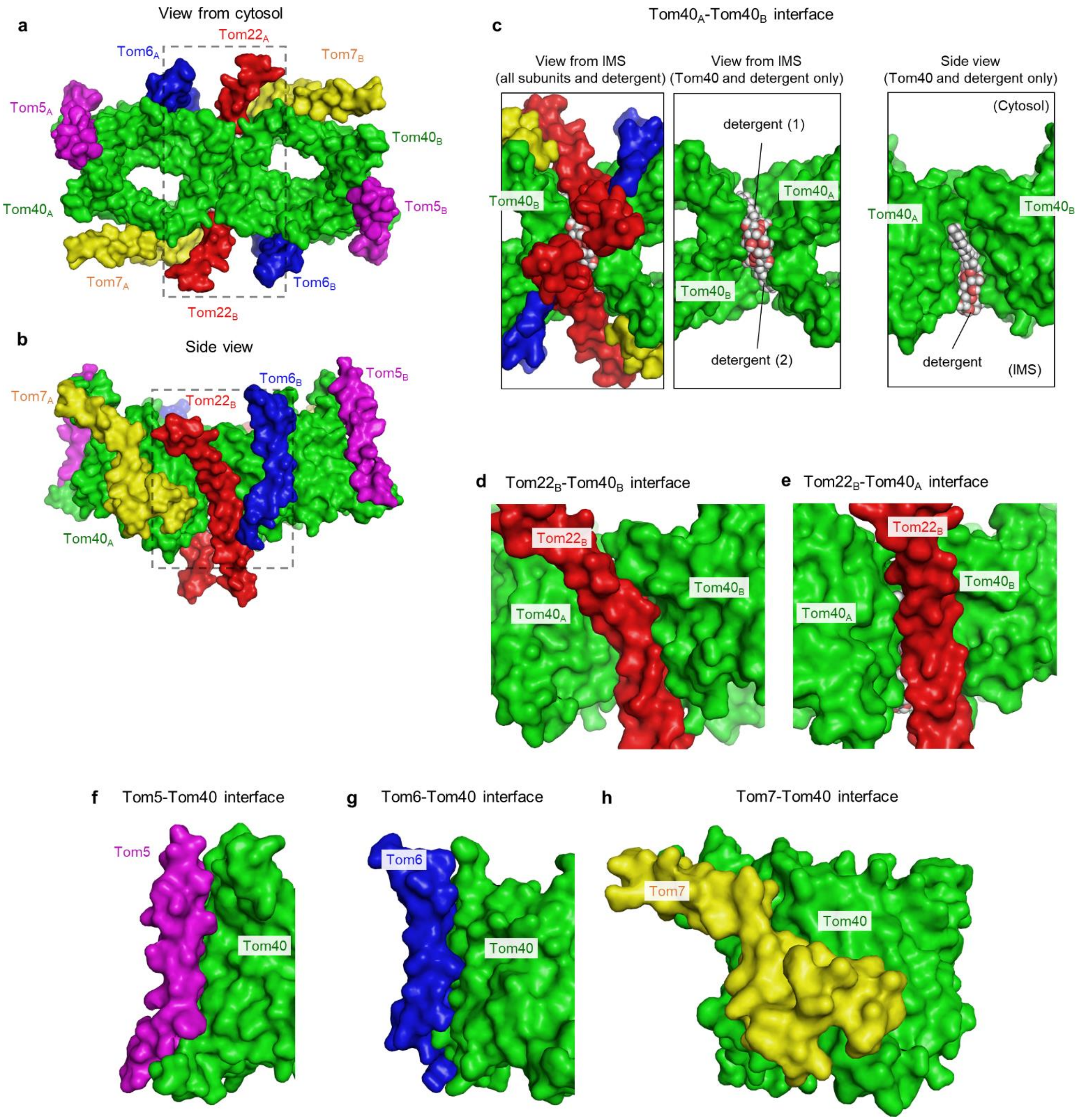


**Supplementary Figure 5. Surface complementarity of Tom subunits at interfaces.**
**a**, **b**, Overview of the dimeric TOM complex in solvent-accessible surface representation. Shown are a view from cytosol (**a**) and a side view (**b**). The color scheme is the same as in Fig. 1. The regions marked by a dashed line are magnified in **c** (with a 180° rotation) and **d**, respectively. **c**, Interface between the two Tom40 subunits. Left, a view from IMS (showing all subunits). Middle, as in the left panel but showing only Tom40 and DDM detergent molecules. Right, as in the middle panel but showing a side view. **d**, **e**, Side views showing the Tom40 and Tom22 interfaces within the same asymmetric unit (**d**) and between the two asymmetric units (**e**). The view directions are the same as in Fig. 2 **a** and **b**, respectively. **f–h**, Side view showing interfaces between Tom40 and other small Tom subunits. The viewing angles are the same as in Fig. 2 **c–e**, respectively.

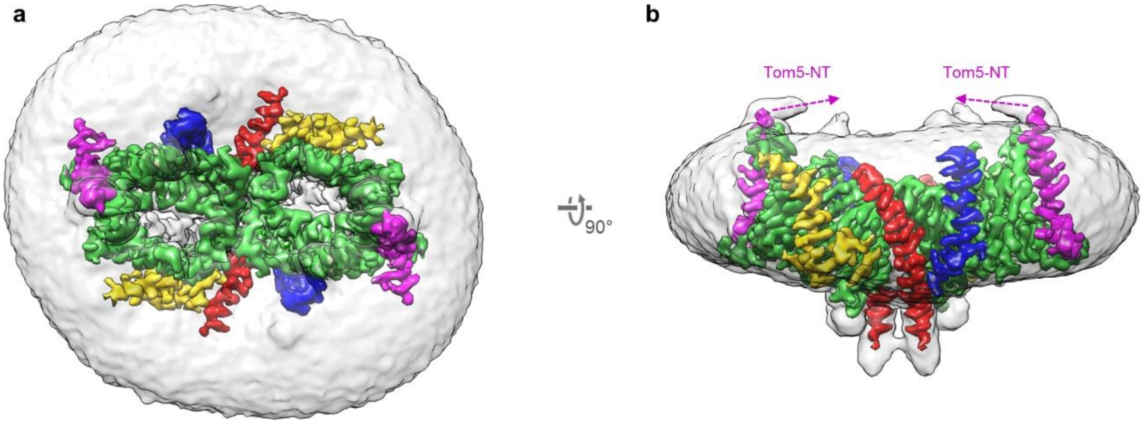


**Supplementary Figure 6. Low-resolution features in the TOM-pALDH complex density map.**
The cryo-EM density map of the TOM-pALDH complex shown at two different contour levels to show strong protein features (colored contour; sharpened and lowpass-filtered at 3.06 Å) and weak features such as the detergent micelle (semitransparent grey contour; unsharpened and lowpass-filtered at 5 Å). The color scheme is the same as in Fig. 1. Shown are a view from cytosol (**a**) and a side view (**b**) with cytosol facing upwards. The magenta dashed arrows indicate approximate directions of the unmodelled N-terminal segments (positions 1–12; Tom5-NT) of the Tom5 subunits, based on weak density features.

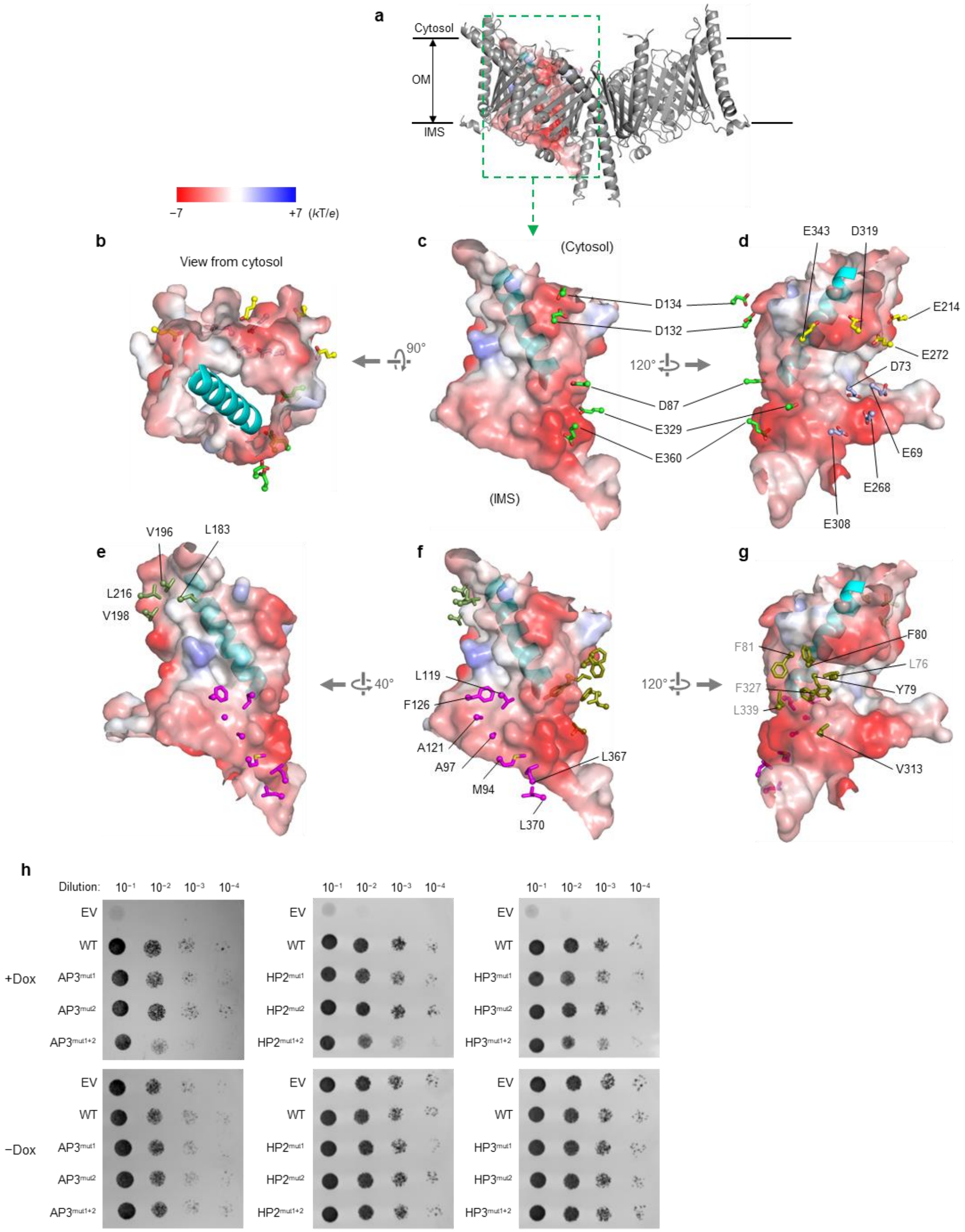


**Supplementary Figure 7. Acidic and hydrophobic patches on the Tom40 pore surface.**
**a**, Overview (side view) of the dimeric TOM complex (grey ribbons) and the Tom40 pore cavity (surface representation; shown for only one Tom40 subunit). OM, outer membrane. **b**–**d**, Surface electrostatics is shown as a heat map overlaid on the pore cavity shown in surface representation. Side chains of acidic amino acids are shown in stick representation (AP1, AP2, and AP3 are in yellow, green, and blue, respectively). In **c**, only AP2 side chains are shown for clarity. The cyan ribbon represents a model for pALDH peptide. **e**–**g**, As in **b**–**d**, but side chains of hydrophobic patches are shown in stick representation (HP1, HP2, are HP3 are in olive, green, and magenta, respectively). Note that some hydrophobic side chains in HP1 (labelled in grey; F81, F327, and L76) are only partially exposed as they are involved in interactions between the α2 segment and the β sheets. **h**, As in Fig. 3i, but with mutants of AP3, HP2, and HP3. $AP3^{mut1}$= E268N/E308N; $AP3^{mut2}$=E69N/D73N; $HP2^{mut1}$=L183S/L216S; $HP2^{mut2}$=V196N/V198S; $HP3^{mut1}$=L119S/A121N/F126N; $HP3^{mut2}$=M94N/A97S. Dox, doxycycline.

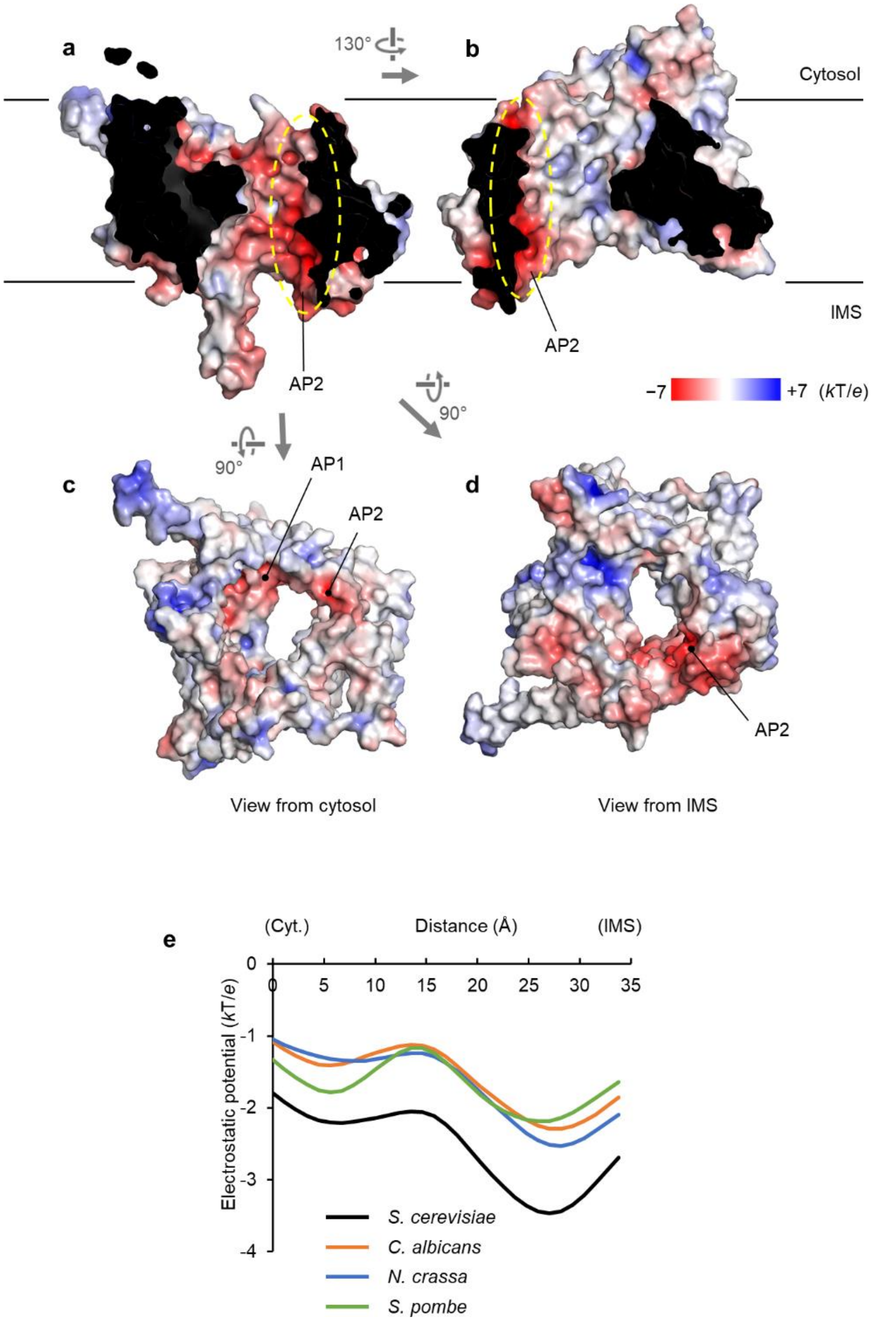


**Supplementary Figure 8. Pore electrostatics of the TOM complex from other fungal species.**
**a**–**d**, As in Fig. 3 **a**–**d**, but with *N. crassa* TOM complex. An *N. crassa* homology model was generated by SWISS-MODEL using the *S. cerevisiae* structure as a template, and electrostatic potential was calculated by Adaptive Poisson-Boltzmann Solver (APBS). Note that unlike the *S. cerevisiae* TOM complex AP3 is not prominent in *N. crassa*. The dashed arrow indicates the likely translocation path. **e**, Electrostatic potential along the translocation path. Homology models for the TOM complexes from indicated species were generated by SWISS-MODEL. After calculating electrostatic potential by APBS, values along the translocation path (dashed arrow in **a**) were extracted and plotted.

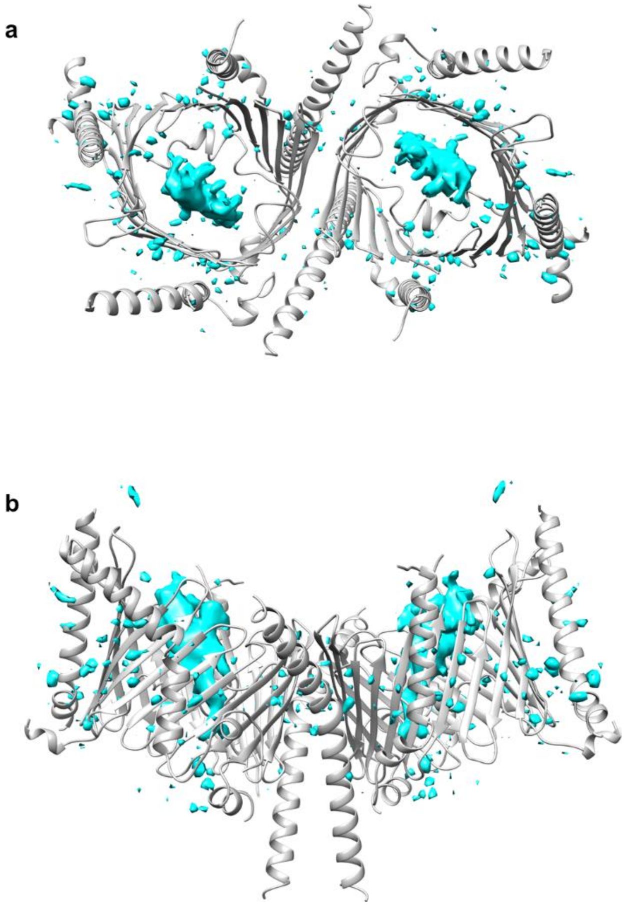


**Supplementary Figure 9. Difference map between the TOM-pALDH and apo TOM reconstructions.**
The difference map subtracting the apo TOM map from the TOM-pALDH map is shown in cyan with the superimposition of the TOM complex model (grey). Small densities overlapping with Tom proteins are mostly due to minor differences in map features that originate from the resolution difference. Shown are a view from cytosol (**a**) and a side view (**b**) as in Fig. 4 **a** and **b**.

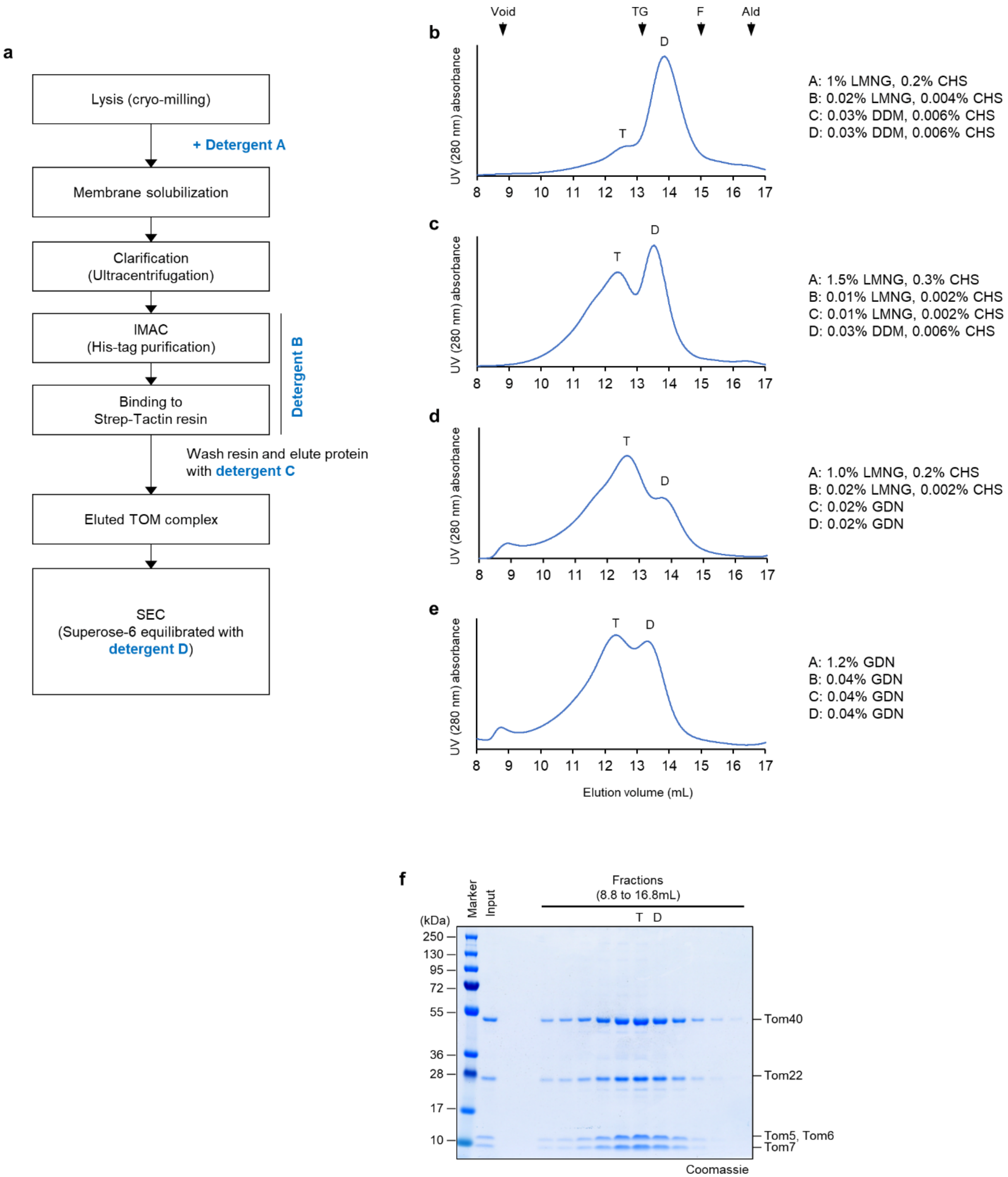


**Supplementary Figure 10. Effects of detergent on the oligomeric state of the TOM complex.**
**a**, Schematic diagram of the TOM complex purification procedure. Different detergent conditions (indicated by blue texts) were explored (see **b**–**e** for specific conditions). **b**–**e**, Detailed SEC profiles of the purified TOM complex purified under different detergent conditions. "D" indicates the dimer peak, and "T" indicates the peak of a higher-order oligomer (tetramer). Positions of the void peak (Void) and peaks of molecular weight standards are indicated by arrowheads. TG, thyroglobulin (670 kDa). F, ferritin (440 kDa). Ald, aldolase (156 kDa). Note that **b**–**d** is the same as in Fig. 5 **a**–**c**, and **b** is the same experiment shown in Supplementary Fig. 1**a**. **f**, SDS-PAGE analysis of peak fractions from the SEC purification shown in **c**. The peak positions are marked with "T" and "D". The SDS gel was stained by Coomassie.

a

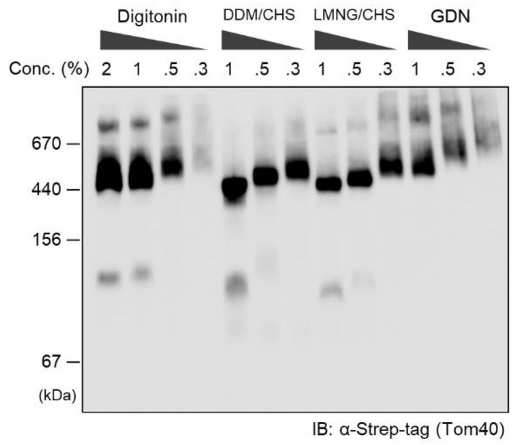


b

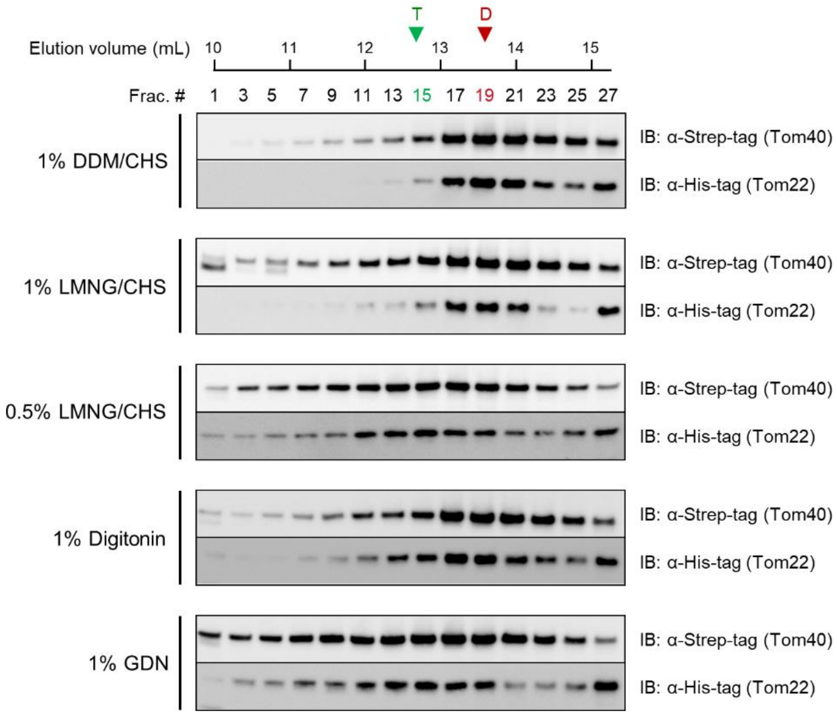


**Supplementary Figure 11. Evaluation of the TOM complex's oligomeric state in crude lysates by BN-PAGE and SEC analysis.**

**a**, Yeast cells overexpressing the TOM complex were lysed, and membranes were solubilized with indicated detergent. The lysates were subjected to BN-PAGE, followed by immunoblotting using an anti-Strep-tag antibody (detecting Tom40-Strep). A gradual decrease of mobility of the TOM complex accompanied by lowered detergent concentrations is likely due to an increased detergent micelle size. **b**, As in **a**, but lysates were injected to a Superose 6 column. The fractions were analyzed by SDS-PAGE and immunoblotting. The column was equilibrated with buffer containing the same detergent used for membrane solubilization at a low concentration (0.03% DDM/0.006% CHS, 0.02% LMNG/0.004% CHS, 0.08% digitonin, or 0.02% GDN). Approximate peak positions are marked with "T" and "D" based on the UV absorbance profiles shown in Supplementary Fig. 9 **b**–**e**. Note that the anti-Strep-tag antibody appears to have substantially lower detection limit (higher sensitivity) than anti-His-tag antibody.

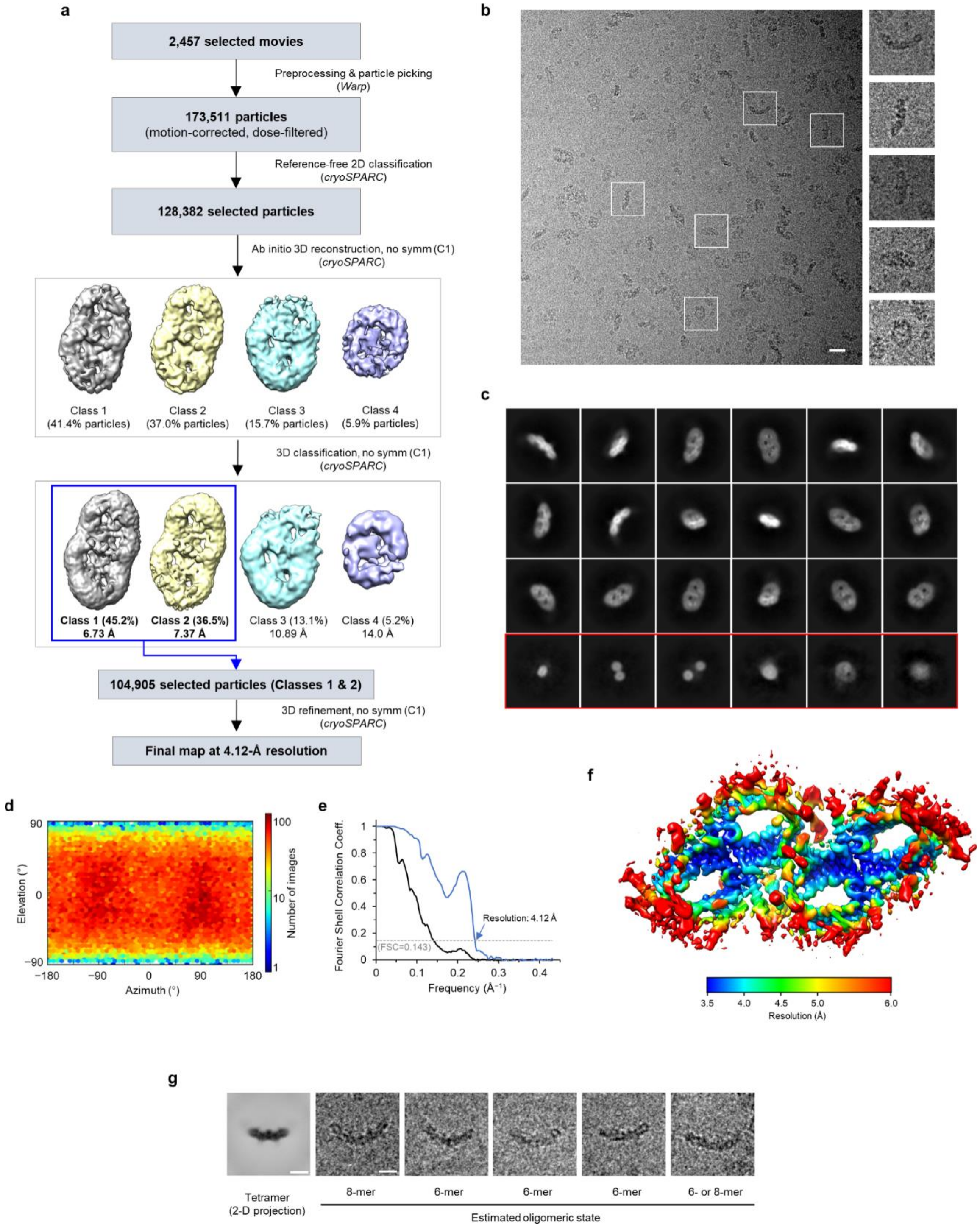


**Supplementary Figure 12. Single-particle cryo-EM analysis of the tetrameric TOM complex.**
**a**, Summary of single-particle image analysis procedure. **b**, A representative motion-corrected micrograph. Scale bar, 20 nm. Right panels show magnified images of selected particles outlined with white squares. The image dimensions are 414 Å (width) by 414 Å (height). **c**, Selected class averages from 2D classification (the second step in **a**). The box dimensions are 460 Å (width) by 460 Å (height). Classes in the red box represent empty micelles and thus excluded from further analysis. **d**, Particle orientation distribution. **e**, Fourier shell correlation (FSC) of the two independently-refined half maps. Blue line, corrected masked FSC. Solid black line, unmasked FSC. **f**, Local resolution map (overlaid on the unsharpened, unfiltered summed map). **g**, Example images of particles larger than the tetramer. The leftmost image shows a 2D projection (side view with the longest width) of the 3D reconstruction of the tetrameric TOM complex. The other images show examples of large particles on micrographs. Estimated oligomeric states are indicated. Scale bar, 100 Å.

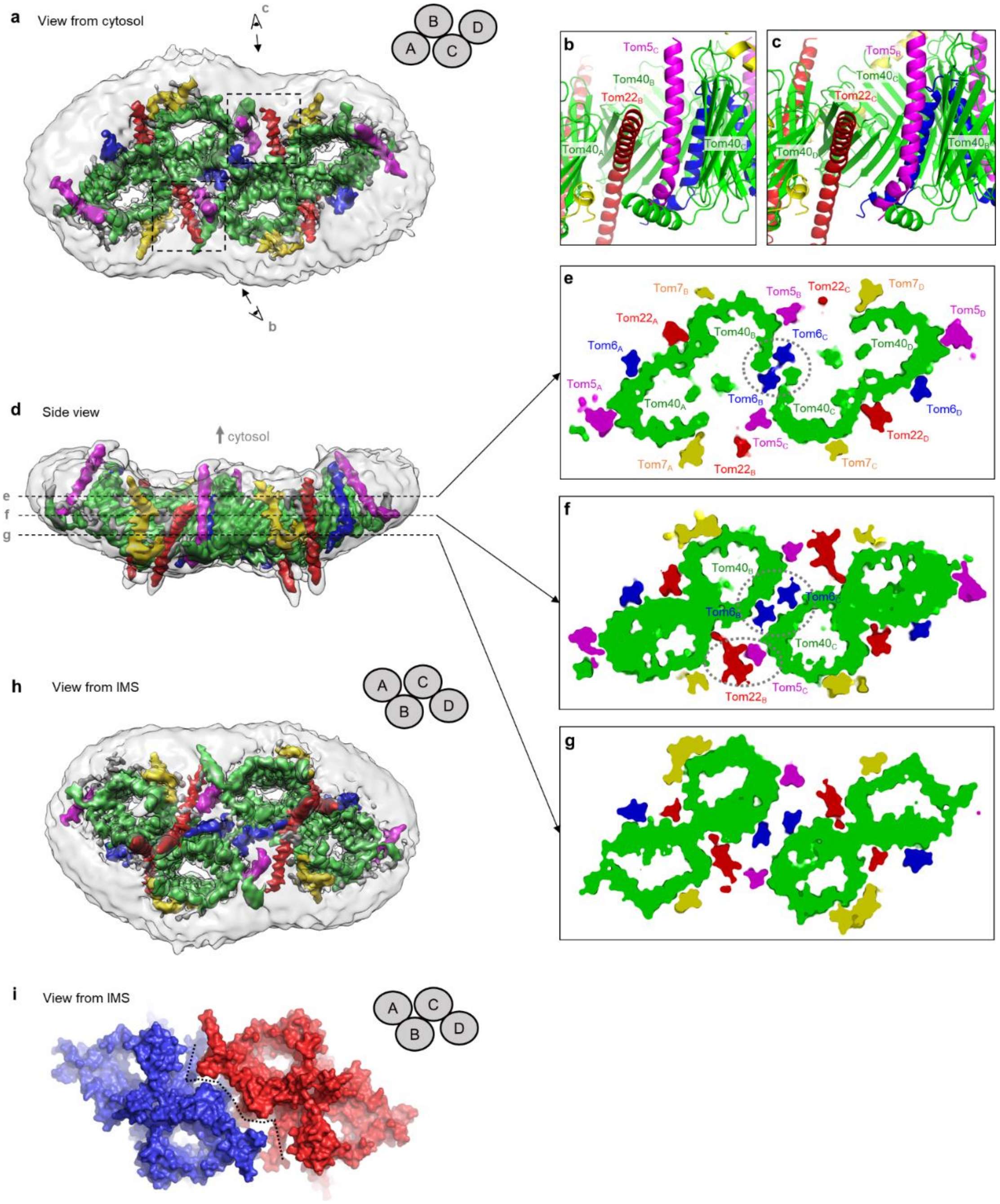


**Supplementary Figure 13. Dimer-dimer interface in the tetrameric TOM complex.**
**a**, Overview (cytosolic view) of the tetrameric TOM complex. The 4.1-Å resolution 3D reconstruction was represented with a composite map showing two different contour levels to show the protein features (colored contour; lowpass-filtered at 4.1 Å) and the detergent micelle (semitransparent grey contour; lowpass-filtered according to local resolution values). Organization of monomeric units are schematized in the upper right corner. Areas marked by dashed rectangles are shown in **b** and **c** (after rotating for a side view) with arrows and eye symbols indicating the viewing directions. **b**, **c**, Side views showing the dimer-dimer contacts between units B and C. Note that the tetramer is not symmetric and that there is a sizeable gap between Tom5$_B$ and Tom22$_C$ (**c**) in contrast to Tom5$_C$ and Tom22$_B$ (**b**). **d**, As in **a**, but showing a side view. Dashed lines indicate cross-sectional planes for cutaway views shown in **e**–**g**. **e**–**g**, Cutaway views (views from cytosol) at different positions along the membrane axis. In **e** and **f**, major interactions mediating the tetramerization are indicated by dashed ovals. Note that in g, there is a gap along the interface (also see **h** and **i**). **h**, As in **a** and **d**, but showing a view from IMS. **i**, Solvent-accessible surface of the tetrameric TOM complex. The dashed line indicates the interfacial gap. The two dimers (A/B and C/D) are in blue and red, respectively.

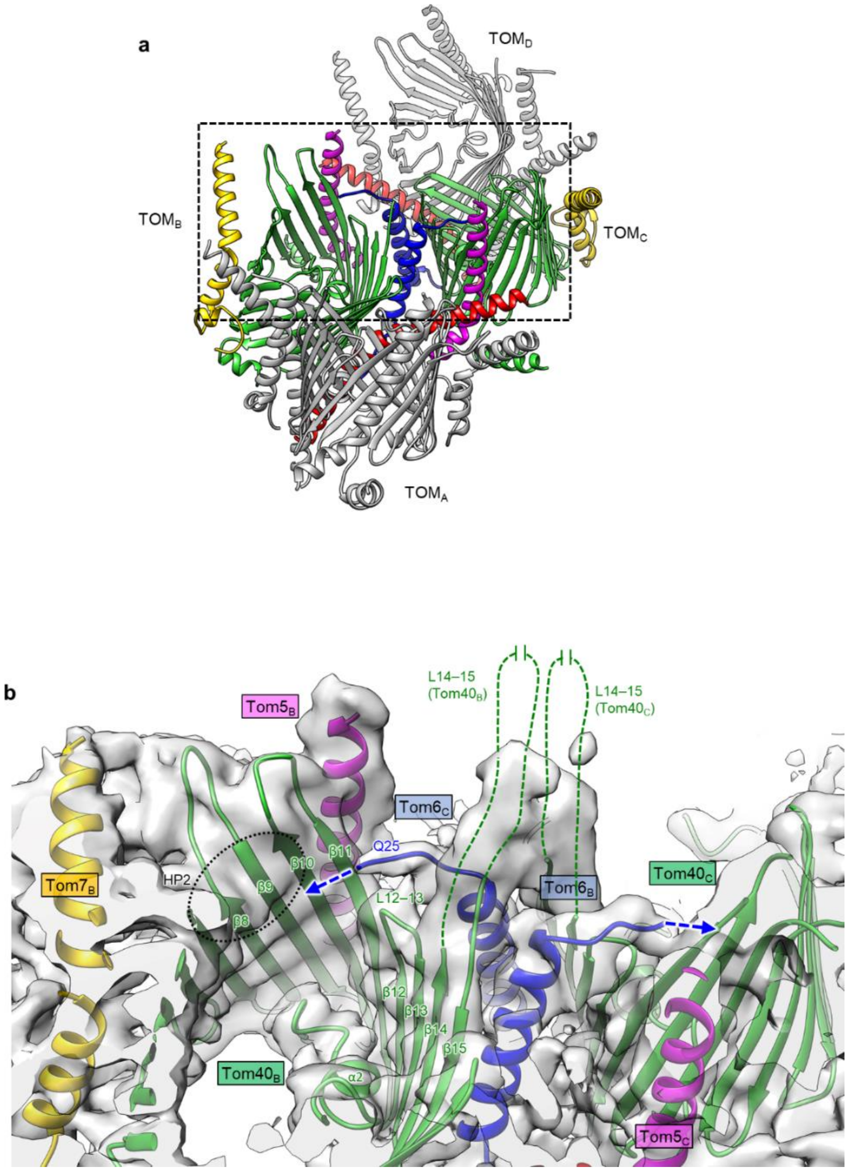


**Supplementary Figure 14. N-terminal segment of Tom6 directed into the Tom40 pore interior.**
**a**, Overview (angled cytosolic view) of the tetrameric TOM complex. Monomeric units B and C are shown in color, and A and D are in grey. The region in the black dashed box is magnified and shown in **b**. **b**, The Cryo-EM density map (semitransparent grey) and the atomic model (in the same color scheme as in Fig. 5) are shown for the B–C interface. The blue dashed arrows indicate the directions of the unmodelled N-terminal segments (residues 1–24) of the $Tom6_C$ and $Tom6_B$ subunits. The black dotted oval indicates the hydrophobic patch HP2. The green dashed lines indicate the unmodeled loop (L14-15; residues 277–294) between β14 and β15 of Tom40.

**Supplementary Table 1. Conservation of inter-subunit polar interactions in fungal TOM complexes**

| *S. cerevisiae* | | | Type of interaction | Other species | | | | |
|---|---|---|---|---|---|---|---|---|
| Position in α-helical Tom subunit | | Position in Tom40 | | Kp | Ca | Sp | Af | Nc |
| Tom22 | W100 | K113 | cation-π | W91–P116 | W96–A117 | W93–P83 | W97–P93 | W87–P93 |
| Tom22 | T105 | H346 | H-bonding | **S96–H346** | **S101–H346** | **S98–Q345** | **S102–Q318** | **S92–H315** |
| Tom22 | S116 | S312 | H-bonding | **S107–S312** | A112–A311 | **S109–S276** | A113–S284 | A103-S280 |
| Tom22 | E120 | R310 | electrostatic | **E111–R310** | **E116–R309** | E113–A274 | **E117–R282** | **D107–R278** |
| Tom22 | E127 | R310 | H-bonding, electrostatic | **E118–R310** | **E123–R309** | E120–A274 | **E124–R282** | **E114–R278** |
| Tom5 | W36 | R52 | cation-π | **W35–R55** | **W34–R56** | L35–K22 | **Y36–R31** | **Y37–R33** |
| Tom6 | T34 | G299 (N) | Weak electrostatic | **Q28–G299*** | **T29–G298*** | **S21–G263*** | **T34–G271*** | **S34–G267*** |
| Tom6 | N38 | G299 (O) | H-binding | **N32–G299*** | **Q33–G298*** | **S25–G263*** | **S38–G271*** | **S38–G267*** |
| Tom6 | I49 (O) | R261 | H-bonding | **I43*–R261** | **I44*–R262** | **L36*–R230** | **L49*–R239** | **L49*–R237** |
| Tom6 | Q50 | W243 | NH-π | **Q44–W243** | **Q45–W244** | **K37–W211** | **H50–W221** | S50–W219 |
| Tom6 | D55 | R261 | H-bonding, electrostatic | **D49–R261** | **D50–R262** | **N42–R230** | **E55–R239** | **E55–K237** |
| Tom6 | L57 (O) | Y307 | H-bonding | **L51*–Y307** | **L52*–Y306** | **L44*–Y271** | **L57*–Y279** | **L57*–Y275** |
| Tom7 | H25 | T163 | H-bonding | **T22–T166** | **K30–S167** | **K19–N134** | R20–G144 | **R20–D142** |
| Tom7 | H29 | L135 (O) | H-bonding | **H26–F138*** | **H34–L139*** | **H23–G105*** | **H24–G115*** | **H24–G115*** |
| Tom7 | N52 | K90 | H-bonding | **N49–K93** | **N57–K94** | R46–K60 | K47–K69 | R47–K71 |
| Tom7 | N52 (O) | K90 | H-bonding | **N49*–K93** | **N57*–K94** | **R46*–K60** | **K47*–K69** | **R47*–K71** |
| Tom7 | L54 (O) | H102 | H-bonding | **L51*–H105** | **L59*–H106** | **I48*–H72** | **F49*–H81** | **L49*–H83** |
| Tom7 | S55 (O) | K90 | H-bonding | **S52*–K93** | **S60*–K94** | **N49*–K60** | **S50*–K69** | **S50*–K71** |
| Tom7 | P56 (O) | T361 | H-bonding | **P53*–S361** | P61*–A360 | P50*–L325 | P51*–I333 | P51*–A330 |
| Tom7 | L57 (O) | K90 | H-bonding | **L54*–K93** | **F62*–K94** | **L51*–K60** | **L52*–K69** | **L52*–K71** |

(Note 1) Species names (% amino acid sequence identity of Tom40 to *S. cervisiae* Tom40): Kp, *Komagataella phaffii* (61.7%); Ca, *Candida albicans* (61.1%); Sp, *Schizosaccharomyces pombe* (42.1%); Af, *Aspergillus fumigatus* (40.1%); Nc, *Neurospora crassa* (38.4%)

(Note 2) "O" and "N" in parentheses indicate interactions mediated by main-chain oxygen and nitrogen atoms, respectively. These interactions may not be amino acid-specific (indicated by asterisk).

(Note 3) Bold print indicates predicted analogous interactions.

**Supplementary Table 2. Cryo-EM image process and atomic model refinement**

| | TOM-pALDH (dimer) | *apo* dimeric TOM | *apo* tetrameric TOM |
|---|---|---|---|
| **Cryo-EM data acquisition and single-particle analysis** | | | |
| Data acquisition | | | |
| Microscope | Titan Krios | Talos Arctica | Titan Krios |
| Acceleration voltage | 300 kV | 200 kV | 300 kV |
| Camera (recording mode) | K2 Summit + GIF (super-resolution mode) | K2 Summit (super-resolution mode) | K2 Summit + GIF (super-resolution mode) |
| Magnification | 43,478x | 43,103x | 43,478x |
| Physical pixel size (Å) | 1.15 | 1.16 | 1.15 |
| Electron dose rate ($e^-$/ Å$^2$/frame) | 1.22 | 1.25 | 1.22 |
| Frame rate (s/frame) | 0.2 | 0.2 | 0.2 |
| Total electron dose ($e^-$/Å$^2$) | 61 | 42.5 (used frames: 1–34) | 43.9 (used frames: 1–36) |
| Defocus range (μm) | −0.8 to −2.5 | −0.9 to −2.5 | −0.9 to −3.0 |
| Number of micrographs collected | 1,766 | 1,528 | 3,104 |
| Number of micrographs used | 1,587 | 976 | 2,457 |
| Image processing and reconstruction | | | |
| Number of extracted particles | 460,148 | 247,202 | 173,511 |
| Box size (pixels) | 256 | 256 | 400 |
| No. of particles in reconstruction | 160,577 | 103,127 | 104,905 |
| Symmetry used for reconstruction | C2 | C2 | C1 |
| Resolution, unmasked (Å) | 4.33 (0.5 FSC)<br>3.93 (0.143 FSC) | 7.1 (0.5 FSC)<br>4.1 (0.143 FSC) | 10.8 (0.5 FSC)<br>7.1 (0.143 FSC) |
| Resolution, masked, corrected (Å) | 3.42 (0.5 FSC)<br>3.06 (0.143 FSC) | 3.90 (0.5 FSC)<br>3.53 (0.143 FSC) | 4.36 (0.5 FSC)<br>4.12 (0.143 FSC) |
| Estimated B-factor (Å$^2$) (cryoSPARC) | 99.7 | 89.0 | 60.8 |
| **Model Refinement (Phenix)** | | | |
| Map pixel size (Å) | 1.15 | 1.16 | 1.15 |
| Map sharpening B-factor (Å$^2$) | −60 | −50 | −60 |
| Map lowpass filter (Å) | 3.06 | 3.53 | 4.12 |
| Refinement resolution limit (Å) | 3.06 | 3.53 | 4.12 |
| Number of atoms, protein | 14,632 (incl. hydrogen) | 7,378 | 14,891 |
| Number of atoms, non-protein | 2,172 (DDM molecules) | 70 (two DDM molecules) | 92 (two DMPC molecules) |
| **Model Statistics** | | | |
| Average B-factor (Å$^2$) | 63.81 | 39.67 | 131.83 |
| r.m.s deviations, bond length (Å) | 0.008 | 0.006 | 0.005 |
| r.m.s deviations, bond angle (°) | 1.25 | 1.22 | 1.17 |
| Ramachandran Plot | | | |
| Favored (%) | 97.09 | 96.34 | 97.20 |
| Outliers (%) | 0.00 | 0.00 | 0.00 |
| Rotamers | | | |
| Favored (%) | 96.62 | 96.74 | 97.11 |
| Outliers (%) | 0.13 | 0.00 | 0.00 |
| MolProbity scores | | | |
| Clash score / percentile | 2.74 / 98% | 4.66 / 95% | 3.83 / 96% |
| Overall score / percentile | 1.22 / 99% | 1.48 / 96% | 1.33 / 98% |
